# OSBP-mediated PI(4)P-cholesterol exchange at endoplasmic reticulum-secretory granule contact sites controls insulin secretion

**DOI:** 10.1101/2023.02.22.529485

**Authors:** Styliani Panagiotou, Kia Wee Tan, Phuoc My Nguyen, Andreas Müller, Affiong Ika Oqua, Alejandra Tomas, Anna Wendt, Lena Eliasson, Anders Tengholm, Michele Solimena, Olof Idevall-Hagren

## Abstract

Insulin secretion is the process whereby insulin-containing granules fuse with the plasma membrane of pancreatic β-cells. Exocytosis is preceded by cargo loading and granule biogenesis at the Golgi, followed by maturation and transport of the secretory granules; processes that require modification of both the protein and lipid composition of the granules. Here, we show that insulin-containing secretory granules form physical contacts with the endoplasmic reticulum. The lipid exchange protein OSBP dynamically redistributes to ER-SG contacts in a process regulated by Ca^2+^ and cytosolic pH, and contributes to cholesterol loading of the granules. This process depends on granular PI(4)P and ER-localized VAPs, and is positively regulated by granular PI4-kinases and negatively regulated by granule-localized Sac2. Loss of Sac2 results in excess accumulation of cholesterol on insulin granules that is normalized when OSBP expression is reduced, and both acute inhibition and siRNA-mediated knockdown of OSBP suppresses glucose-stimulated insulin secretion without affecting insulin production or intracellular Ca2+ signaling. In conclusion, we show that lipid exchange at ER-granule contact sites is involved in the exocytic process, and propose that these contacts act as reaction centers with multimodal functions during insulin granule maturation.

## INTRODUCTION

Blood glucose homeostasis critically depends on the appropriate release of insulin from pancreatic β-cells. Insulin is produced as a pro-peptide that is packaged into secretory granules that bud from the trans-Golgi network. These immature granules undergo a series of maturation steps that involve changes in the composition of both granular lipids and proteins and that coincide with the maturation of insulin (Omar-Hmeadi and Idevall-Hagren, 2021). A typical β-cell contains between 5000-15000 secretory granules (SGs) (Müller et al., 2021; Sato Tamiko, 1981), but only around 50-100 of these have the ability to fuse with the plasma membrane (Barg et al., 2002; Olofsson et al., 2002). These granules belong to the readily releasable pool and are situated at the plasma membrane where they await a triggering Ca^2+^ signal. Prolonged stimulation of insulin secretion will deplete this pool and new SGs must be recruited from the large reserve pool to sustain secretion. This recruitment primarily involves newly synthesized SGs and depends on granule transport along microtubules as well as acquisition of specific priming and docking factors, and interactions with components of the secretory machinery, including t-SNARE family members and voltage-dependent Ca^2+^ channels (Barg et al., 2002; Barg et al., 2010; Bokvist et al., 1995; Bracey et al., 2020; Hoboth et al., 2015; Ivanova et al., 2013). Defects in the replenishment of release-competent SGs have been observed in type-2 diabetes and likely contribute to disease development (Gandasi et al., 2018).

The mechanisms by which SGs dock and fuse with the plasma membrane are well-characterized and to a large extent similar to those of other neuro-endocrine cells. However, the steps preceding the arrival of granules at the plasma membrane are largely unclear. These steps include the action of numerous small GTPases, including Rab3, Rab27 and Rab2, and the corresponding Rab effectors, but it is not well understood how these reactions are regulated (Omar-Hmeadi and Idevall-Hagren, 2021). Progression through the endo-lysosomal system requires the cooperative action of Rabs and phosphoinositides, which together controls the recruitment of both phosphoinositide-metabolizing enzymes and Rab effectors and ensures unidirectionality throughout the membrane trafficking cascade (Di Paolo and De Camilli, 2006). To what extent the same principal guide progression through the regulated secretory pathway is not known. The trans-Golgi network is rich in phosphatidylinositol 4-phosphate (PI(4)P), and this lipid is directly involved in the formation of nascent SGs via recruitment of the GTPase Arf1 (Cruz-Garcia et al., 2013; Dittié et al., 1996). The membrane of the newly formed SGs likely contains high concentrations of PI(4)P, both as a consequence of its Golgi origin and through the action of PI4-kinases (Olsen et al., 2003). We recently described the involvement of the PI(4)P phosphatase Sac2 in insulin SG maturation and exocytosis (Nguyen et al., 2019). Lack of Sac2 resulted in accumulation of PI(4)P on the SG surface and in an inability of granules to stably dock and fuse with the plasma membrane. This mechanism resembles vesicle maturation in yeast cells, where the ER-localized PI(4)P phosphatase Sac1 is recruited to the vesicle surface (Ling et al., 2014). In these cells, recruitment occurs through an interaction with the lipid transport protein Osh4p, which also directly contributes to PI(4)P removal by exchanging it for cholesterol (de Saint-Jean et al., 2011; Ling et al., 2014). To what extent a similar mechanism operates during regulated secretion of insulin is not known. However, observations that loss of the mammalian Osh4p homologue OSBP leads to impaired insulin secretion and that SG cholesterol accumulation is required for insulin release are consistent with the existence of such a mechanism (Hao et al., 2007; Hussain et al., 2018; Xia et al., 2008).

Here, we now show that loss of Sac2 in β-cells results in excess accumulation of cholesterol on the surface of insulin SGs through an OSBP-dependent process. Recruitment of OSBP to the granule surface is controlled by granular PI(4)P and dynamically regulated by cytosolic Ca^2+^ and pH, with knockdown or inhibition of OSBP resulting in impaired insulin secretion. These results unravel a novel SG maturation pathway in β-cells and also demonstrate for the first time a role of ER-SG contact sites in regulated secretion.

## MATERIALS AND METHODS

### Reagents and plasmids

TopFluor Cholesterol (23-[dipyrrometheneboron difluoride]-24-norcholesterol was from Avanti Polar Lipids, Inc. Cholesterol Assay Kit (Cell-Based) was from Abcam (Ab122116). OSW-1 was from Cayman Chemical. Cal-590-AM and Cal520-AM were from AAT Bioquest. BCECF was from Life technologies. Plasmids pLJM1-FLAG-GFP-OSBP was from Addgene (Plasmid #134659; (Lim et al., 2019)), GFP-Sac2 and VAPA-GFP were gifts from Pietro De Camilli (Yale University, New Haven, CT; (Nakatsu et al., 2015)) GFP-Rab3a, NPY-mCherry, NPY-mNG and VAMP2-pHluorin were gifts from Sebastian Barg, Uppsala University, Sweden. RA-Sec61β and GB-Dcp1 were gifts from Gia Voeltz (Addgene plasmid #153978, #153979). Halo-OSBP was created by amplifying OSBP from pLJM1-FLAG-GFP-OSBP (Addgene #134659) and inserting it by Gibson Assembly (New England Biolabs) into the BglII/NotI sites of a customized cloning vector bearing an N-terminal Halo-tag (pENTR20-Halo-C1). This vector was recombined using Gateway LR reaction (Invitrogen) into a customized Gateway-enabled pCDH vector (System Biosciences) and expressed under control of a EF1α promoter. GB-Rab3 was created by amplifying mouse Rab3a ORF and inserting it into the XhoI/SacII site of the GB-Dcp1 using the following primers: FW: 5’-GATCTCGAGTACCGGTTGGATCAGGATCAATGGCCTCAGCCACAGAC-3’ RV: 5’-GGGCCCGCGGTCAGCAGGCGCAATCC-3’. GB-OSBP was generated by amplifying OSBP from Halo-OSBP using the following primers: FW: 5’-CAAGTCCGGACTCAGATCCATGGCTGCTACGGAGC-3’, RV: 5’-CAGTTATCTAGATCCGGTGGATCTCAGAAAATGTCCGGGCATG-3’ followed by ligation to pGB-Dcp1 using Xho1/Xma1. GB-CAAX was generated by releasing Rab3 from GB-Rab3 with Nhe1/Age1 followed by ligation to pCIBN-CAAX (Idevall-Hagren et al., 2012). RA-CAAX was generated by releasing RA from RA-Sec61β with Nhe1/Age1 followed by ligation to pGB-CAAX. GB-E-Syt3 was generated by releasing E-Syt3 from pEGFP-E-Syt3 (Giordano et al., 2013) with Nhe1/Xho1 followed by ligation to pGB-Dcp1. E-Syt3-GB was generated by releasing E-Syt3 from pE-Syt3-EGFP (Giordano et al., 2013) with Nhe1/Age1 followed by ligation to pTOM20-GB (Xie et al., 2022). RA-Rab3 was generated by releasing Rab3 from GB-Rab3 with Xho1/SacII followed by ligation to pRA-Sec61β (Lee et al., 2020). GB-Sec61β was generated by releasing GB from pGB-Dcp1 (Lee et al., 2020) with Nhe1/Xho1 followed by ligation to pRA-Sec61β. Adenoviral particles (E5 serotype) carrying RA-Sec61β and GB-Rab3a under the control of CMV-promoters were produced by Vector Biolabs (Malvern, PA). The following antibodies were used in the study: OSBP polyclonal (catalogue no. PA5-30110, host: rabbit, 1:300, Invitrogen), VAPA polyclonal (HPA009174, Atlas Antibodies, host: Rabbit, 1:400), VAPB polyclonal (HPA013144, Atlas Antibodies, host: Rabbit, 1:400), PI4K2A monoclonal (sc-390026, Santa Cruz, host: mouse, 1:100), β-actin monoclonal (clone C4, sc-47778, Santa Cruz Biotechnology, host: mouse, 1:300), IP3R1 polyclonal (catalogue no. ab264281, Abcam, host: rabbit, 1:500), Rab3 monoclonal (catalogue no. 107 111, Synaptic Systems, host: mouse, 1:500), Chromogranin A polyclonal (catalogue no. 10529-1-AP, Proteintech, host: Rabbit, 1:400), insulin polyclonal (catalogue no. A0564, Dako, host: guinea pig, 1:1000), calnexin (PA5-34754, Invitrogen, host: Rabbit, 1:500), GolgB1 (BS-13356R, Bioss, host: rabbit, 1:500).

### Cells and transfection

MIN6 cells were used between passages 18 to 35 and kept in cell culture medium composed of DMEM supplemented with 4.5 g/L glucose, 2 mM L-glutamine, 100 U/mL Penicillin, 100 μg/mL Streptomycin, 50 μM 2-mercaptoethanol, and 15% fetal bovine serum (all from Life Technologies). The cells were cultured in a humidified atmosphere at 37°C and 5% CO_2_. Reverse transfections were carried out in 100 μL of OptiMEM with 0.5 μL Lipofectamine 2000 or Lipofectamin 3000 (all from Life Technologies), 0.1-0.4 μg plasmid DNA, and 200,000 cells. The transfection was terminated after 3-5 hrs, and imaging was done between 22 and 30 hrs after transfection. INS-1 832/3 cells were cultured in RPMI-1640 with 11 mM D-glucose, supplemented with 10% FBS, 10 mM HEPES, 1 mM sodium pyruvate, 50 μM β-mercaptoethanol, and 1% penicillin/streptomycin in a 37°C/5% CO_2_ incubator.

### Viral transduction of MIN6 cells

MIN6 cells were infected with 10 µl of high titration virus in 100 μL of culture medium (RA-Sec61b; 5.5×10^10^ PFU/mL, GB-Rab3; 4.5×10^10^ PFU/mL; Vector Biolabs, Malvern, PA). After 3 hrs, the coverslips were washed with medium, and the cells were cultured for at least two days to allow the expression of the contact sites reporter before performing experiments.

### Pancreatic islet isolation and culture

All animal experimental procedures were approved by the local ethics committee for use of laboratory animals in Uppsala, Sweden. Islets of Langerhans were isolated from 4–12-month-old C57Bl6J mice by collagenase digestion of the pancreas followed by hand-picking of individual islets. After isolation, the islets were cultured for 1-3 days in RPMI 1640 medium containing 5.5 mM glucose, 10% fetal calf serum, 100 U/mL penicillin and 100 µg/mL streptomycin at 37°C in a 5% CO_2_ humidified atmosphere. 20-50 freshly isolated islets were infected with 10 µL of high titration virus in 100 μL of culture medium (RA-Sec61b; 5.5×10^10^ PFU/mL, GB-Rab3; 4.5×10^10^ PFU/mL; Vector Biolabs, Malvern, PA). After 3 hrs, the coverslips were washed with medium and the cells were cultured for at least two days to allow the expression of the contact sites reporter before performing experiments. Human islets were isolated from the pancreas of cadaveric organ donors by the Nordic Network for Clinical Islet Transplantation Uppsala (ethical approval by Uppsala Regional Ethics Board) with written donor and family consent for use in research. Work with human tissue complied with all relevant ethical regulations for use of human tissue in research and the study was approved by the Uppsala Regional Ethics Board (2006/348). Isolated islets were cultured free-floating in sterile dishes in CMRL 1066 culture medium containing 5.5 mM glucose, 10% fetal calf serum (FCS), 2 mM l-glutamine, streptomycin (100 U/ml), and penicillin (100 U/ml) at 37 °C in an atmosphere of 5% CO_2_ for up to 2 weeks.

### siRNA-mediated knockdown

Sac2 knockdown was performed with siRNA targeting mouse gene sequence Inpp5f: 5’-GGAAUGCGGUAUAAACGAATT-3’ and was from Ambion (Life Technologies). OSBP knockdown was performed using ON-TARGET plus SMART pool Mouse OSBP siRNA from Dharmacon. MIN6 cells were transfected in 12 well-plates with siRNA using Lipofectamine 2000, followed by a second round of transfection 3-4 hours later using RNAiMax (Life Technologies) according to the manufacturer’s instructions with final siRNA concentration of 50 nM and 20 nM for Sac2 and OSBP, respectively. ON-TARGET plus NON-targeting Pool from Dharmacon was used as control. INS-1 832/3 cells were transfected with 40 nM of Control (Ambion Silencer Select Negative Control #1 siRNA; Cat no: 4390843) or VAP-A+B siRNA (Ambion Silencer Select Pre-Designed siRNAs; Cat#:4390771), using Lipofectamine 2000 according to the manufacturer’s instructions.

### Quantatitive RT-PCR

Sac2 knockdown was confirmed by quantitative RT-PCR using QuantiTect SYBR Green RT-PCR kit (Qiagen) and the following primers: Sac2-fwd: 5’TAAGGAGAGCCAGAGAAGCCA-3’, Sac2-rev: 5-CAGCAGCACTTCCACATCTCT-3’; GAPDH-fwd: 5’-ACTCCACTCACGGCAAATTC-3’; GAPDH-rev: 5’-TCTCCATGGTGGTGAAGACA-3’. PCR reactions were performed using Light Cycler 2.0 (Roche). Results are expressed as ΔΔCt, normalized to the expression in control samples.

### Western blot

Cells were lysed on ice with RIPA buffer (50 mM Tris-HCl, pH 7.4, 1% NP-40, 0.5% Na-deoxycholate, 0.1% SDS, and 10% EDTA) supplemented with a protease inhibitor cocktail (Roche). Samples were cleared by centrifugation and protein concentration was measure by the

Bradford assay. OSBP knockdown was confirmed by Western blotting using OSBP Polyclonal Antibody (1:1000, Invitrogen) and β-actin mouse monoclonal (clone C4, sc-47778, 1:300, Santa Cruz Biotechnology).

### Immunofluorescence

Cells grown on monolayers were fixed in 4% paraformaldehyde in either PBS or PHEM buffer (60mM PIPES, 25mM HEPES, 10mM EGTA, 4mM MgSO_4_, pH 6.9) for 20 mins at RT, followed by 3 washes of buffer. Permeabilization was performed using 0.1-0.2% Triton X-100 for 10-15 mins, followed by 3 washes of buffer, and blocking for 1 hr in buffer containing 2% BSA. Primary antibody incubation was carried out overnight at 4°C in blocking buffer, washed 3 times and incubated for 1 hr with secondary antibody in blocking buffer. Coverslips were then washed 3 times, mounted in Prolong Glass (Invitrogen) and left to dry for at least 18 hrs before imaging. Where NPY-mNeonGreen was visualized in fixed samples, the secondary antibody incubation included 5 nM of Fluotag-X2-anti-mNeonGreen conjugated to Atto488 (Nanotag Biotechnologies). For INS-1 832/3 cells, permeabilization was carried out with 0.5% (v/v) Triton X-100/PBS for 10 mins, followed by blocking with 1% BSA/PBS for 1 hr and incubation with primary antibodies in 0.1% BSA/PBS at 4°C overnight. Cells were then washed in PBS and incubated in secondary antibodies for 1 hr before being washed and mounted with ProLong Diamond Antifade Mountant with DAPI (ThermoFisher).

### Cholesterol visualization

TopFluor cholesterol was diluted (0.6 mg/mL) in methanol:water (95:5 vol%) followed by sonication. The dissolved cholesterol was transferred to glass tubes and dried under N_2_, followed by resuspension in 4 mg/mL fatty acid free BSA. MIN6 cells were loaded with 1 μM TopFluor cholesterol and incubated for 3 hrs followed by additional culture in a cholesterol-free medium for 15-18 hrs prior live cell imaging. Filipin cholesterol staining was performed following the manufacturer’s instructions (Ab122116 - Cholesterol Assay Kit, Abcam) and imaging was performed 0-4 hrs after staining.

### Measurements of intracellular Ca^2+^ and pH

To measure changes in the intracellular Ca^2+^ concentrations, MIN6 cells or mouse islets were preincubated at 37°C for 30 mins in an experimental buffer (see below) supplemented with 2 µM of the AM-ester-form of the Ca^2+^-indicator Cal-520. The cells were subsequently washed twice with experimental buffer before visualization by fluorescence microscopy (excitation 491 nm, emission 530/50 nm). Intracellular pH was determined using the ratiometric dye BCECF. MIN6 cells were incubated with 2 µM of the AM-ester-form of BCECF for 30 mins at 37°C, followed by two washing steps to remove residual indicator and visualization by fluorescence microscopy (excitation 442 nm and 491 nm, emission 530/50 nm).

### Insulin secretion measurements

MIN6 cells were cultured in 12-well plates to approximately 70% confluency. Cells were washed three times with PBS, pre-incubated in experimental buffer (125 mM NaCl, 4.9 mM KCl, 1.2 mM MgCl_2_, 1.3 mM CaCl_2_, 25 mM Hepes, and 0.1% BSA) with 3 mM Glucose for 30 mins at 37°C. Cells were then incubated for 30 mins in experimental buffer with 3 mM glucose, 20 mM glucose or 3 mM glucose and 30 mM KCl. The supernatants were collected for insulin measurement using a Mouse Insulin ELISA kit (Mercodia, Uppsala, Sweden). Isolated mouse islets of Langerhans (8-10 islets per experiment) were placed in a closed 10 µL chamber made of Teflon tubing with a fine mesh plastic net at the outlet that prevented the islets from escaping. Islets were subsequently perfused with experimental buffer. The perfusate was collected in 5 min fractions and insulin content in the fractions was determined using a mouse insulin ELISA kit (Mercodia). Cells were subsequently collected, diluted in acidic ethanol (75% EtOH and 15% HCl) and sonicated for 2×10 sec to determine the insulin content.

### OSW-1 treatment

Cells were incubated in experimental buffer containing either 0.1% DMSO or 20 nM OSW-1 (dissolved in DMSO) (Toronto Research Chemicals) for 30 mins at 37°C before imaging or ELISA sample collection. INS-1 832/3 cells were incubated with or without 50 nM OSW-1 at 37°C for 15 min and fixed with 4% PFA in PBS for 20 min before immunofluorescence.

### Proximity ligation assay

PLA was performed according to the manufacturer’s recommendation (Navinci™ NaveniFlex MR). Primary antibodies against VAP-A (HPA009174, rabbit polyclonal, Sigma Aldrich), SEC61 (ab15576, rabbit polyclonal, Abcam) and Rab3 (catalog no. 107 111, mouse monoclonal, Synaptic Systems) were used at 1:200 dilution.

### Fluorescence microscopy

All experiments, unless otherwise stated, were performed at 37°C in an experimental buffer containing 125 mM NaCl, 4.9 mM KCl, 1.2 mM MgCl_2_, 1.3 mM CaCl_2_, 25 mM Hepes, 3 mM D-Glucose, and 0.1% BSA (pH 7.4). Cells were preincubated in imaging buffer for 30 mins before experiments, and continuously perfused with the same buffer at the stage of the microscope. Where required, Halo-tagged proteins were labeled with 40 nM of Halo-tag ligand bearing JFX646 (Grimm et al., 2021) for 20 mins in experimental buffer and unbound ligand was washed out for 10 mins in the same buffer. Confocal microscopy was performed on a Nikon Eclipse Ti-2 equipped with a Yokogawa CSU-10 spinning disc confocal unit and a 100×/1.49-NA plan Apochromat objective (Nikon) as previously described (Idevall-Hagren et al., 2013). Briefly, excitation light was provided by 491-nm, 561-nm DPSS lasers and 640-nm diode laser (all from Cobolt), and fluorescence was detected through interference filters 530/50-nm (GFP, mNG, Alexa488), 590/20-nm (mCherry, RA, Alexa568) or 650LP filter (JF646) with a back-illuminated EMCCD camera (DU-888; Andor Technology) controlled by MetaFluor software (Molecular Devices). TIRF microscopy was performed on two different setups. A custom-built prism-type TIRF microscope equipped a 16×/0.8-NA water-immersion objective (Nikon) was used for low-magnification imaging of cell populations as previously described (Idevall-Hagren et al., 2013). It is built around an E600FN upright microscope (Nikon) contained in an acrylic glass box thermostated at 37°C by an air stream incubator. DPSS lasers (Cobolt) provided 491-nm and 561-nm light for excitation of GFP and mCherry. The laser beams were merged with dichroic mirrors (Chroma Technology), homogenized, and expanded by a rotating Light Shaping Diffuser (Physical Optics) before being refocused through a modified quartz dove prism (Axicon) with a 70° angle to achieve total internal reflection. Laser lines were selected with interference filters (Semrock) in a motorized filter wheel equipped with a shutter (Sutter Instruments) blocking the beam between image captures. Fluorescence from the cells was detected at 530/50 nm for GFP (Semrock interference filters) or 597LP for mCherry (Melles Griot glass filter) with a EM-CCD camera (Andor DU-887) under MetaFluor software control. TIRF imaging was also performed on an inverted Nikon Ti-E equipped with a TIRF illuminator and a 60X 1.45-NA objective (all Nikon). Excitation light was provided by diode-pumped solid-state lasers (491-nm, 561-nm) or diode lasers (640-nm; all from Cobolt). Lasers were merged with dichroic mirrors (Chroma technologies) and excitation light was selected with bandpass filters mounted in a filter wheel (Sutter Instruments Lambda 10-3) and delivered to a fiber optic cable and delivered to the TIRF illuminator. Excitation light was reflected through the objective with a dichroic mirror (ZET405/488/561/640m-TRFv2, Chroma Technologies) and emission light was separated using interference (530/50 nm, 590/20-nm) or longpass (650LP) filters (Semrock) mounted in a filter wheel (Sutter instruments Lambda 10-3). An electronic shutter was used to block light exposure between image captures. Emission light was detected using an Orca-AR camera controlled by MetaFluor software (Molecular Devices Corp.). Filipin imaging was visualized on a laser scanning confocal microscope (Zeiss LSM 780) equipped with a Plan-Apo 20X/0.8-NA objective using a 405-nm diode laser. Confocal images from INS-1 832/3 cells were captured with a Zeiss LSM-780 inverted microscope from the Facility for Imaging by Light Microscopy at Imperial College London, UK using a 63x/1.40 Oil DIC Plan-Apochromat objective.

### Electron microscopy

Mouse or human islets were hand-picked under a microscope and groups of ∼50 isolated islets were fixed in Millonig’s buffer with 2.5% glutaraldehyde overnight, post-fixed in 1.0% osmium tetroxide, dehydrated and embedded in AGAR 100 (Oxfors Instruments Nordiska AB, Sweden). Before visualization, 70–90 nm sections were cut, mounted, and contrasted. The samples were examined in a JEM 1230 electron microscope (JEOL-USA, Inc, MA, USA).

### 3D segmentation of ER-insulin SG contact sites

Segmentation of ER-SG contact sites was performed on focused ion beam electron microscopy (FIB-SEM) volumes of isolated mouse pancreatic islets previously acquired for and available via the OpenOrganelle platform (https://openorganelle.janelia.org/) under a CC BY 4.0 license (Xu et al., 2021). Small volumes with contact sites were cut in FIJI (Schindelin et al., 2012) and manual segmentation of ER and SG was performed with the Labkit FIJI plugin (Arzt et al., 2022). Ribosomes were segmented by black-and-white-thresholding in Microscopy Image Browser (Belevich et al., 2016). Segmentation results were rendered with ORS Dragonfly (https://www.theobjects.com/dragonfly/index.html).

### Image analysis

TIRF microscopy images were analyzed offline using ImageJ. Briefly, for analysis of VAMP2-pHluorin and Cal-520 experiments, cell footprints were manually identified and regions of interest covering the edges of the adherent cells were drawn. Intensity changes within these regions during the experiments were measured and exported to Excel. All data points were background corrected, followed by normalization to the pre-stimulatory level (F/F0). Analysis of enrichment of tagged proteins to insulin SGs was performed using a pipeline assembled in Cell Profiler. Insulin SGs were segmented using the NPY channel. Cells positive for the protein of interest were segmented and all insulin SGs not in an OSBP-positive cell were discarded. The median fluorescence in a 2-pixel wide ring around each insulin SG was defined as the background and the enrichment of the protein of interest was calculated by dividing the mean of fluorescence in each insulin granule with its corresponding background. To determine the percentage of insulin SG positive for a protein, we set the enrichment threshold at 2 standard deviations from the mean. For each treatment, the standard deviations for the basal condition were computed and used for their corresponding treatment conditions. Confocal images from INS-1 832/3 cells were analyzed using coloc-2 plugin in Fiji.

### Statistical analysis

Statistical analysis was performed using 2-way ANOVA followed by Tukey’s or Kruskall-Wallis (for non-parametric data) posthoc tests. Where a single test group was compared against a treatment group, we performed Student’s 2-tailed unpaired t-test or Mann-Whitney U-test (for non-parametric data). Paired analysis was performed with Student’s 2-tailed paired t-test. We did not perform a power analysis and the experiments did not have a predetermined sample size, but this was instead chosen based on experience. At least three independent experiments (performed on separate days) were used for all analyses.

## RESULTS

### ER-SG contact sites

Ultrastructural examination of β-cells from mouse pancreatic islets of Langerhans using transmission electron microscopy revealed the presence of ribosome-free ER domains in close proximity to insulin SGs (Fig. 1A). To obtain a more detailed view of these contacts we performed volumetric reconstruction of individual ER-granule contacts in FIB-SEM data of mouse β-cells. We frequently observed ribosome-free ER in close proximity (10-20 nm) to insulin-containing SGs, and in many cases the ER traced the outlines of the granules in a pattern resembling contacts the ER makes with other vesicular organelles (Fig. 1B, C) (Dong et al., 2016; Lim et al., 2019; Mesmin et al., 2013). Proximity ligation assay (PLA) using antibodies against the general ER-marker Sec61β and Rab3 (SG) further demonstrated the existence of physical contacts between the ER and SGs in mouse islet cells (Fig. 1D, E and Suppl. Fig. 1). Next, we repeated the PLA using an antibody against VAP-A, a protein commonly involved in membrane contact site formation, for ER targeting. This resulted in a 4-fold increase in detected PLA puncta (Fig. 1F, G), despite very similar distribution patterns of Sec61β and VAP-A in traditional immunofluorescence staining (Suppl. Fig. 1), indicating that VAP-A may be enriched at the ER-SG interface. We also noted strong PLA signal from islet cells negative for insulin, suggesting that close proximities between ER and SG also exist in the other islet cell types (Fig. 1E, F). Next, we wanted to determine if these contacts existed in live cells and to explore their putative dynamics. We therefore developed a proximity sensor based on dimerization-dependent fluorescent proteins (Alford et al., 2012). This sensor makes use of two quenched monomers, RA and GB, that form a fluorescent heterodimer when brought in proximity to each other. To test the applicability of this method for visualization of membrane contact sites, we designed a version for reporting ER-PM contact sites by targeting RA to the surface of the ER using Sec61β and GB to the plasma membrane through C-terminal lipidation (GB-CaaX). Co-expression of these two constructs resulted in distinct fluorescence that was restricted to the cell periphery and overlapped with an ER marker (Suppl. Fig. 2A). As a positive control, we targeted both RA and GB to the ER, resulting in strong fluorescence (Suppl. Fig. 2B), and as a negative control we expressed either GB-CaaX or RA-Sec61β. When expressed alone, GB-CaaX showed no detectable fluorescence while RA-Sec61β showed very weak fluorescence, consistent with previous work showing incomplete quenching of RA fluorescence (Suppl. Fig. 2C, D) (Alford et al., 2012). To determine at what distance from each other the two moieties must be to reconstitute a fluorescent protein, we fused GB to the N- or C-terminus of ER-localized E-Syt3 and expressed it together with RA-CaaX. Whereas C-terminally localized GB interacted with RA at the plasma membrane, we could not detect any fluorescence signal from cells expressing N-terminally-localized GB (Suppl. Fig. 2E). Based on the structure of E-Syt2 (Schauder et al., 2014), we therefore estimate that RA and GB must be <20 nm from each other, which is a typical inter-membrane distance at membrane contact sites (MCSs) (Scorrano et al., 2019). We next localized GB to the surface of SGs by fusing it to Rab3a and co-expressed it with RA-Sec61β. This resulted in strong punctate fluorescence which overlapped with markers of both the ER (ER-oxGFP) (Fig. 2A-C) and SGs (NPY-mNG) (Fig. 2D, E). Whereas all contacts overlapped with the ER marker, only a subset of SGs showed overlap with the contact site reporter. The overlap with SGs was highest close to the plasma membrane, where most granules were in contact with the ER, while granules located more centrally in the cell exhibited fewer ER contacts (Fig. 2D). This demonstrates the existence of a subpopulation of SGs defined by physical tethering to the ER. Time-lapse imaging revealed that the contacts were dynamic and moved together with SGs, indicating physical tethering, and not just proximity, to the ER.

**Figure 1:**
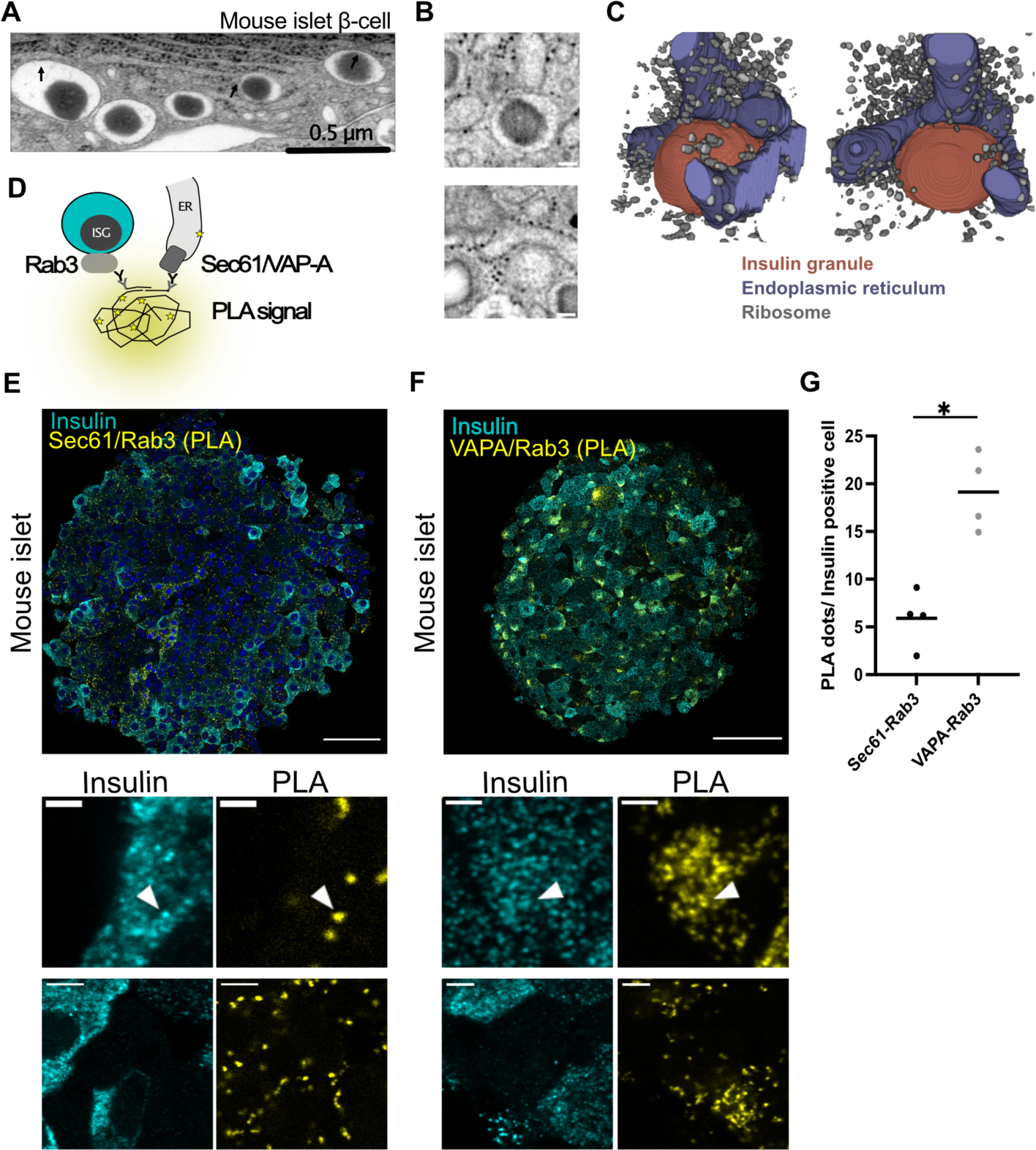
Membrane contact sites between the endoplasmic reticulum and insulin granules. **A.** Transmission electron microscopy image of a mouse islet β-cell showing close proximities between insulin granules and the endoplasmic reticulum (arrows). **B.** x-y slices at different z-positions from FIB-SEM images showing the proximity between an insulin granule and the ER in a mouse β-cell. Scale bar: 100 nm. **C.** Volumetric renderings of B showing the insulin granule in red, the ER in purple and ribosomes in grey. **D.** Principle of Proximity Ligation Assay (PLA) for detection of ER-insulin granule contacts using antibodies against the ER (Sec61/VAP-A) and insulin granules (Rab3a). **E.** Confocal image of a mouse islet with insulin immunostaining shown in cyan and ER-granule contact sites (Sec61b/Rab3a) detected with PLA in yellow. Magnified images below show β-cells (top pair) and non β-cells (bottom pair) containing contact sites. Scale bars: 50/5µm. **F.** Confocal image of a mouse islet with insulin immunostaining shown in cyan and ER-granule contact sites (VAP-A/Rab3a) detected with PLA in yellow. Magnified images below show β-cells (top pair) and non β-cells (bottom pair) containing contact sites. Arrows identify PLA puncta overlapping with insulin granules. Scale bars: 50/5µm. **G.** Mean number of PLA puncta per insulin-positive cell detected using the indicated antibody pairs. (n=4 islets for each antibody pair, P<0.05, unpaired, 2-tailed Student’s t-test).

**Figure 2:**
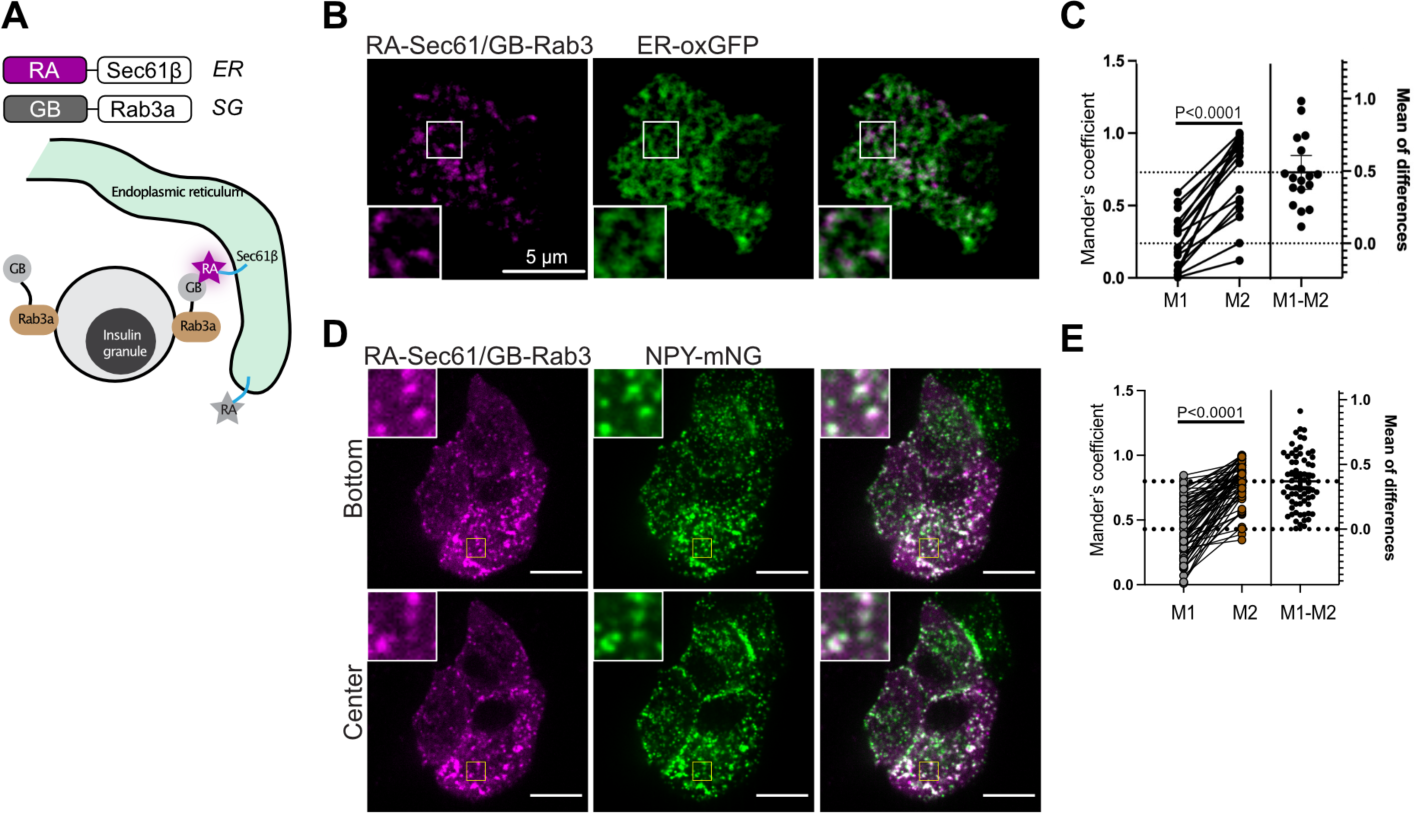
Dimerization-dependent fluorophore reporter for ER-SG contact sites. **A.** Diagram showing the use of conditional fluorophores as a probe for ER-granule contacts. The ER-targeted dimerization-dependent RFP (RA) dimerizes with the granule targeted nonfluorescent booster (GB), revealing ER-SG contact sites. **B.** Confocal microscopy images of a MIN6 cell expressing the ER marker ER-oxGFP (green) and the RA-Sec61b/GB-Rab3 contact site reporter (magenta). Scale bar: 5µm. **C.** Quantification of the colocalization between ER and the contact site reporter (M1) and the contact site reporter and ER (M2) (n=17 cells; 2-tailed Student’s t-test). **D.** Confocal microscopy images of a MIN6 cell expressing the insulin granule marker NPY-mNG (green) and the RA-Sec61b/GB-Rab3 contact site reporter (magenta). Scale bar: 10µm. **E.** Quantification of the colocalization between granules and the contact site reporter (M1) and the contact site reporter and granules (M2) (n=53 cells; 2-tailed Student’s t-test).

### OSBP localizes to ER-SG contact sites in a PI(4)P-dependent manner

MCSs are hubs for lipid exchange reactions, and several lipid transport proteins are found at multiple MCSs. Oxysterol binding protein 1 (OSBP) localizes to MCSs by binding to PI(4)P in target membranes and to VAPs in the ER membrane (Levine and Munro, 2002; Mesmin et al., 2013; Wyles et al., 2002). It is primarily found at the ER-Golgi interface where it exchanges Golgi PI(4)P for ER cholesterol, resulting in removal of PI(4)P through subsequent dephosphorylation by the ER-resident PI(4)P-phosphatase Sac1 (Mesmin et al., 2013; Mesmin et al., 2017). Immunofluorescence imaging of MIN6 β-cells showed that OSBP primarily localized to the Golgi, and to a lesser extent to the plasma membrane and the cytosol (Fig. 3A). Preincubation with the OSBP inhibitor OSW-1 (20 nM, 30 min), which blocks the lipid exchange and stabilizes OSBP at PI(4)P-rich membranes, caused a striking redistribution of OSBP to punctate structures in the cytosol (Fig. 3A, C). A similar pattern was seen in mouse islet β-cells (Fig. 3B, C), and co-immunostaining for insulin revealed that some of these structures were insulin SGs (Fig. 3A, B). Next, we expressed fluorescently tagged OSBP (GFP-OSBP or Halo-OSBP) and the SG marker NPY (NPY-mNG) in MIN6 cells and observed the live cells on a confocal microscope. OSBP had a predominant Golgi localization, although 2-5% of the granules were found in close apposition to OSBP-positive structures (Fig. 3D, E). Strikingly, addition of OSW-1 resulted in rapid enrichment of OSBP on SGs, with around 10-20% of the granules being positive for OSBP after 30 mins incubation with OSW-1 (P<0.0001 *versus* control; Fig. 3D, E). To test if these granules represent a subpopulation in proximity to the ER, we again took advantage of the dimerization-dependent fluorescent protein. We made a version where OSBP was fused to the non-fluorescent GB-subunit and co-expressed this together with RA-Sec61β and a SG marker (NPY-mNG) in MIN6 β-cells. Under resting conditions, there was essentially no red fluorescence overlapping with the location of SGs. Addition of 20 nM OSW-1 resulted in the rapid appearance of fluorescent puncta that overlapped with, or were in close proximity to, SGs, demonstrating that OSBP becomes enriched at locations where SGs are within 20 nm of the ER (Fig. 3F, G)). Since OSBP stabilization at MCSs typically require binding to the ER-localized VAPs, we next tested how OSBP recruitment to SG was affected by siRNA-mediated knockdown of VAP-A and VAP-B in rat insulinoma INS-1 832/3 cells. In control cells, addition of OSW-1 induced the expected accumulation of OSBP at SGs, but this response was completely absent in cells with reduced VAP expression (Fig. 3H). Immunofluorescence staining of VAP-A/B and insulin SG revealed proximity between the two signals, but, in contrast to OSBP, there was no further enrichment of VAPs at SGs following OSW-1 addition (Suppl. Fig. 3A-D). Importantly, the addition of OSW-1 was also without effect on the overall Golgi and ER morphology (Suppl. Fig. 3E-G). Next, we tested if the OSW-1-induced recruitment of OSBP to SGs contributes to ER-SG contact site formation. We expressed the ER-SG proximity reporter together with the SG marker NPY-mNG and exposed the cells to a control buffer or a buffer containing OSW-1. Neither treatment affected the number of SGs in proximity to the ER, suggesting that OSBP is recruited to pre-existing ER-SG contacts (Fig. 3I, J).

**Figure 3:**
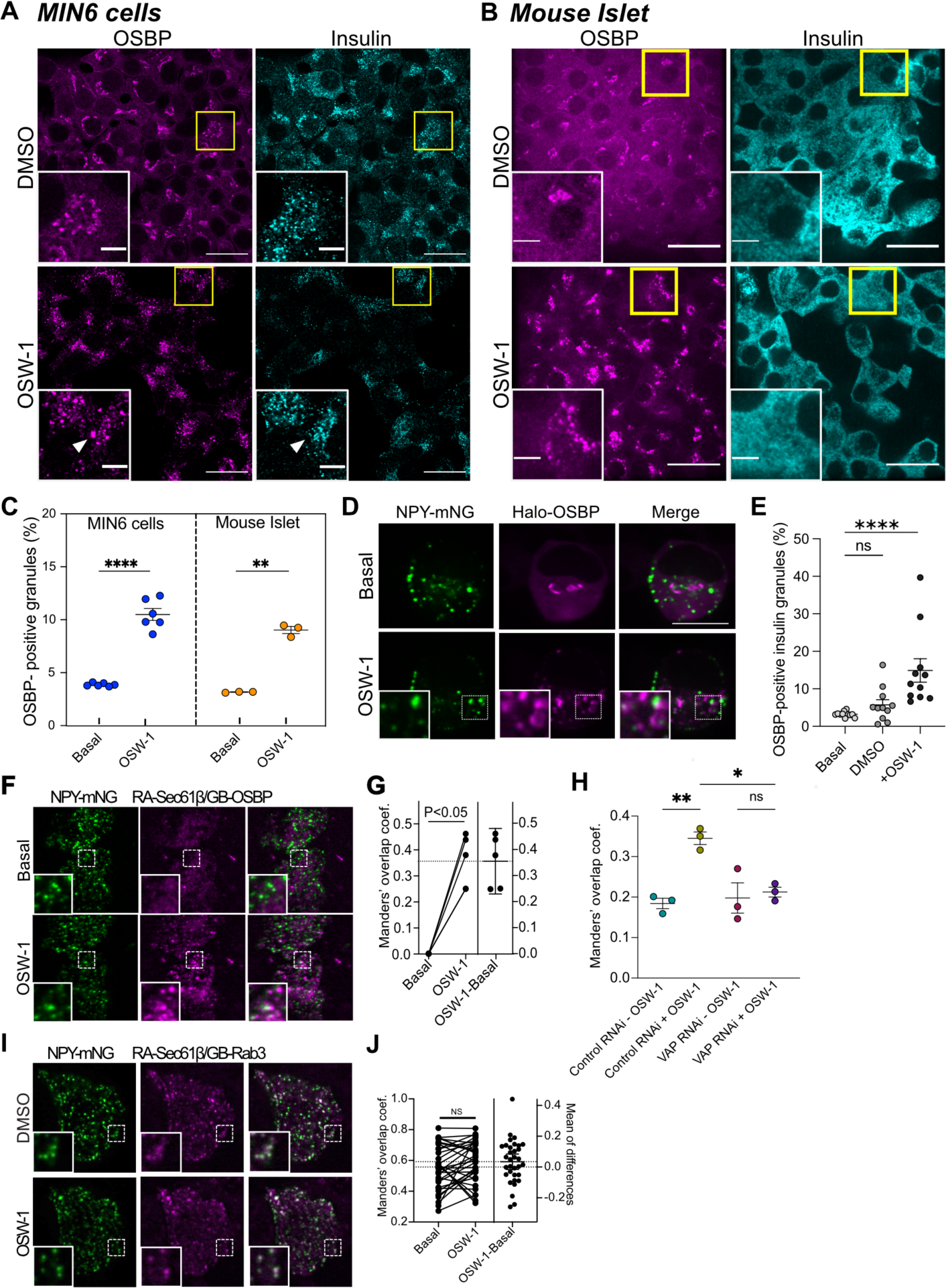
OSBP localizes to insulin granules in proximity to the ER. **A.** Confocal microscopy images of MIN6 cells treated with OSW-1 (20 nM for 20 mins) or vehicle (DMSO) and immunostained for insulin (cyan) and OSBP (magenta). Insets magnified from boxed regions. Arrowheads show an overlapping punctum. Scale bars: 20/5 µm. **B.** Confocal microscopy images of a mouse islet treated with OSW-1 (20 nM for 20 mins) or vehicle (DMSO) and immunostained for insulin (cyan) and OSBP (magenta). Insets magnified from boxed regions. Scale bars: 20/5µm. **C.** Quantification of immunostained OSBP-positive insulin granules in MIN6 cells or mouse islet cells treated with OSW-1 (20 nM for 20 mins) or vehicle (basal). Each dot represents the mean of one experiment (** P<0.01; **** P<0.0001; 2-tailed, unpaired Student’s t-test). **D.** Confocal microscopy images of MIN6 cells expressing NPY-mNG and Halo-OSBP, imaged live under basal conditions or following treatment with OSW-1 (20 nM for 20 mins). Insets magnified from boxed regions. Scale bar: 10 µm **E.** Quantification of OSBP-positive insulin granules from MIN6 cells expressing NPY-mNG and Halo-OSBP imaged live on a confocal microscope and treated with OSW-1 (20 nM for 20 mins) or vehicle (DMSO). **F.** TIRF microscopy images of MIN6 cells expressing NPY-mNG, RA-Sec61b, and GB-OSBP, imaged live under basal conditions and following treatment with OSW-1 (20 nM for 20 mins). Insets magnified from boxed regions. Scale bar: 10 µm. **G.** Manders coefficient of RA-Sec61b/GB-OSBP signal overlapping with NPY-mNG puncta before and after treatment with 20 nM OSW-1. **H.** Quantification of the overlap between OSBP and insulin-containing granules in control and VAP-A/B knockdown INS1 832/3 cells following 30 min incubation with DMSO (-OSW1) or 50 nM OSW-1 (+OSW1). (** P<0.001, * P<0.01; n=30-40 cells from 3 separate experiments; One-way ANOVA). **I.** Confocal microscopy images of a MIN6 cell expressing RA-Sec61b/GB-Rab3 and NPY-mNG under basal conditions (DMSO) and after 20 mins treatment with 20 nM OSW-1. **J.** Manders coefficient of RA-Sec61b/GB-Rab3 signal overlapping with NPY-mNG puncta before and after treatment with 20 nM OSW-1.

OSBP interacts with PI(4)P in target membranes through its C-terminal PH-domain (Levine and Munro, 2002). Consistent with PI(4)P-dependent binding to SGs, we found that the wild-type PH-domain of OSBP, but not that of a PI(4)P-binding-deficient mutant (PH^RR107/108EE^), stabilized at SGs upon addition of OSW-1 (Fig. 4A, B). We previously showed that SGs contain PI4-kinase activity and that this is required for the recruitment of the PI(4)P phosphatase Sac2 (Nguyen et al., 2019). To test if the binding of OSBP to SG depended on granular PI4-kinase activity, we pre-treated cells expressing Halo-OSBP and NPY-mNG with PI4-kinase inhibitors and exposed them to OSW-1. We found that the recruitment of OSBP to SG was prevented by the broad PI4-kinase inhibitor phenylarseneoxide (PAO) but not by the more selective class IIIβ (PIK93 and PI4KI) or class IIIα (GSK-A1) inhibitors (Fig. 4C, D and Suppl. Fig. 4). Immunostainings as well as live cell imaging revealed the presence of class IIα PI4-kinases on insulin SGs, making this a likely candidate for maintaining the granular PI(4)P pool (Fig. 4E-G).

**Figure 4:**
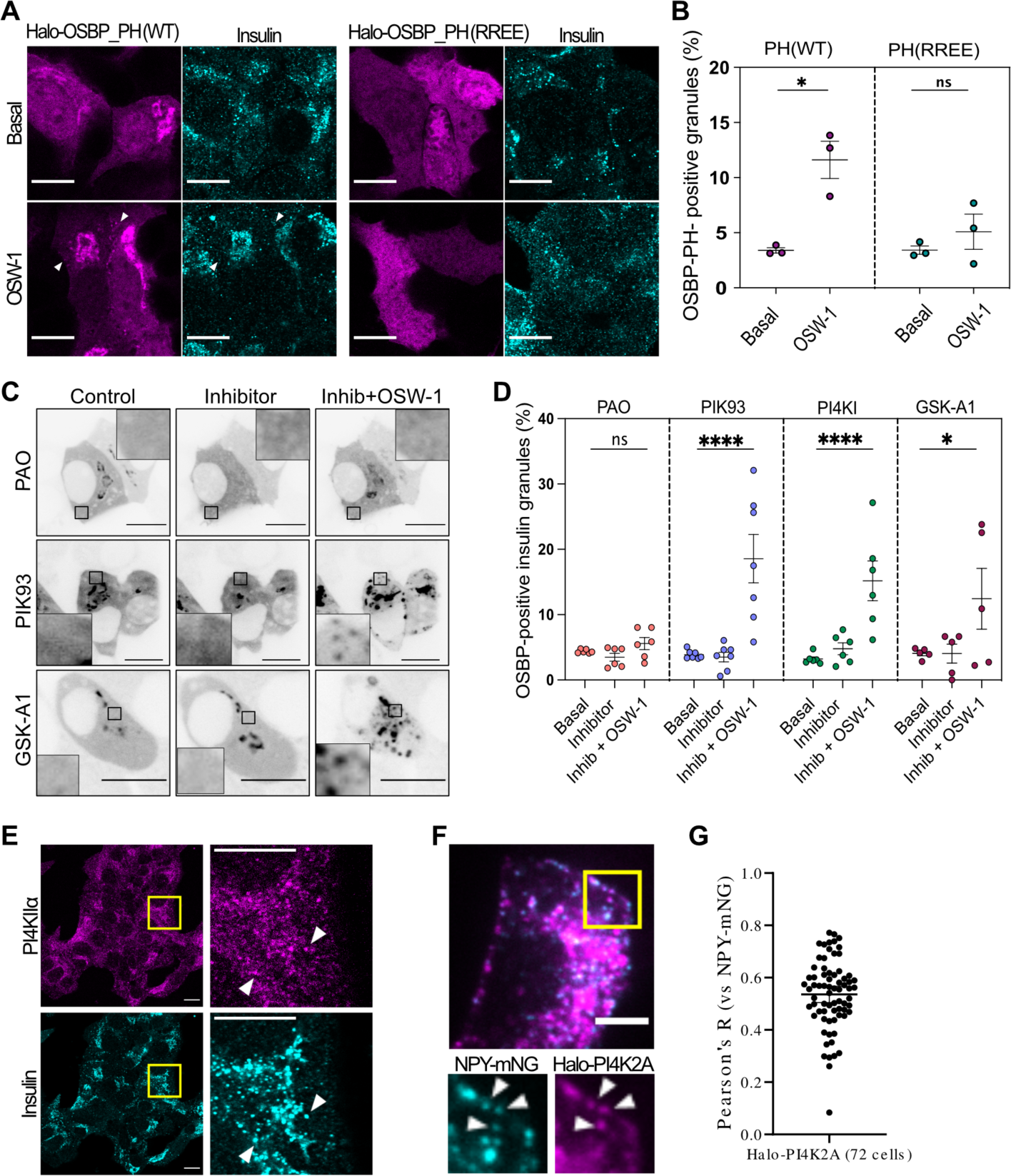
OSBP recognizes a granule-localized pool of PI(4)P. **A.** Confocal microscopy images of MIN6 cells expressing Halo-OSBP-PH or the non-PI4P binding RR107/108EE mutant, treated with 50 nM OSW-1 for 20 mins or vehicle, fixed and immunostained against insulin. **B.** Quantification of OSBP-PH positive insulin granules from experiments done as in A (* P<0.05; 2-tailed unpaired Student’s t-test; n=3). **C.** Confocal images of MIN6 cells expressing NPY-mNG (not shown) and Halo-OSBP, treated with the indicated PI4-kinase inhibitors for 20 mins followed by treatment with 20 nM OSW-1 for 20 mins. The Halo-OSBP channel is shown (inverted), insets magnified from boxed regions. Scale bar: 5 µm. **D.** Quantifications of OSBP-positive granules from experiments performed as in C. Each dot represents the average from one experiment. (* P<0.05; **** P<0.0001; 2-tailed unpaired Student’s t-test; n=5-7 replicates). **E.** Confocal microscopy images of MIN6 cells immunostained against PI4K2A and insulin. Boxed region magnified to the right. Scale bar: X nm. **F.** Confocal images and zoom-ins of MIN6 cells expressing the granule marker NPY-mNG and Halo-PI4K2A. **G.** Pearson’s R for NPY-mNG and Halo-PI4K2A (n=72 cells).

### OSBP binding to SGs is regulated by Ca^2+^ and cytosolic pH

Although constitutive contacts between the ER and SGs are abundant, OSBP does not seem to be a major component of these since it primarily localizes to the Golgi under resting conditions. The addition of OSW-1 induces accumulation of OSBP at SGs, indicating that there is a constitutive weak or transient interaction between OSBP and SGs that can be stabilized by preventing lipid exchange. We next asked whether OSBP enrichment at SGs might be enhanced under certain conditions. The binding of OSBP to the Golgi is counteracted by increases in the cytosolic Ca^2+^ concentration, likely through the inhibition of PI4-kinases at the Golgi with accompanying reduction in Golgi PI(4)P (Balla et al., 2008; Malek et al., 2021). Ca^2+^ is a key second messenger in β-cells, and glucose stimulation causes voltage-dependent Ca^2+^ influx that triggers insulin granule exocytosis. To test if voltage-dependent Ca^2+^ influx had an impact on OSBP distribution, we depolarized MIN6 β-cells expressing GFP-OSBP or Halo-OSBP. We found that Ca^2+^ influx triggered the immediate dissociation of OSBP from the Golgi, and the process was reversed when the depolarizing agent was removed (Fig. 5A, B and Suppl. Fig. 5). TIRF microscopy imaging of MIN6 β-cells expressing OSBP-GFP and the ER-marker Sec61β-mRFP showed the appearance of OSBP puncta after depolarization that partially overlapped with the ER marker (Fig. 5C). The accumulation of OSBP puncta coincided with the rise of Ca^2+^, and there was a linear correlation between the extent of OSBP accumulation and the increase in Ca^2+^ indicator fluorescence (Fig. 5D-F). Similar displacement was also seen in response to Ca^2+^ release from the ER triggered by the addition of carbachol (Suppl. Fig. 5) and in response to an elevation of the glucose concentration from 3 to 20 mM (Fig. 5L-N). These experiments show that Ca^2+^ elevations trigger the release of OSBP from the Golgi, perhaps making it available for interactions at other MCSs. PI(4)P was recently shown to act as a pH sensor through protonation at acidic pH, which results in reduced affinity for effector proteins (Shin et al., 2020). Depolarization of β-cells is associated with acidification of the cytosol (Salgado et al., 1996), and we therefore tested if pH changes may contribute to OSBP displacement from the Golgi. To accomplish rapid pH changes, we exposed MIN6 cells loaded with the pH-sensitive dye BCECF to 10 mM NH_4_Cl. This caused a rapid alkalinization, seen as an increase in fluorescence. Washout of NH_4_Cl resulted in transient acidification that was followed by pH-normalization at the pre-stimulatory level within 1 min (Suppl. Fig. 5). Subsequently, we exposed MIN6 cells expressing OSBP-GFP and loaded with the Ca^2+^ indicator Cal590 to 10 mM NH_4_Cl. The acute alkalization was without effect on both cytosolic Ca^2+^ and OSBP localization, while washout of NH_4_Cl caused dramatic dissociation of OSBP from the Golgi without accompanying change in cytosolic Ca^2+^ (Fig. 5G, H). Importantly, alkalinization of the cytosol with 10 mM NH_4_Cl completely prevented the depolarization-induced dissociation of OSBP from the Golgi while only slightly suppressing the rise of cytosolic Ca^2+^ (Fig. 5G, J and Suppl. Fig. 5). Next, we co-expressed OSBP-GFP and the SG marker NPY-mCherry in MIN6 cells and imaged them with TIRF microscopy. Depolarization resulted in an enrichment of OSBP at SGs which was counteracted by alkalinization of the cytosol with NH_4_Cl, indicating that Ca^2+^ increases facilitate the interaction between OSBP and granules at least partly through acidification of the cytosol (Fig. 5H-K). Importantly, we found that acute addition of the OSBP inhibitor OSW-1 was associated with an acidification of the cytosol, seen as a drop in fluorescence intensity of cytosolically expressed pHluorin, implicating either roles of OSBP in pH regulation or off-target effects of the compound that may contribute to the redistribution of the protein (Suppl. Fig. 5).

**Figure 5.**
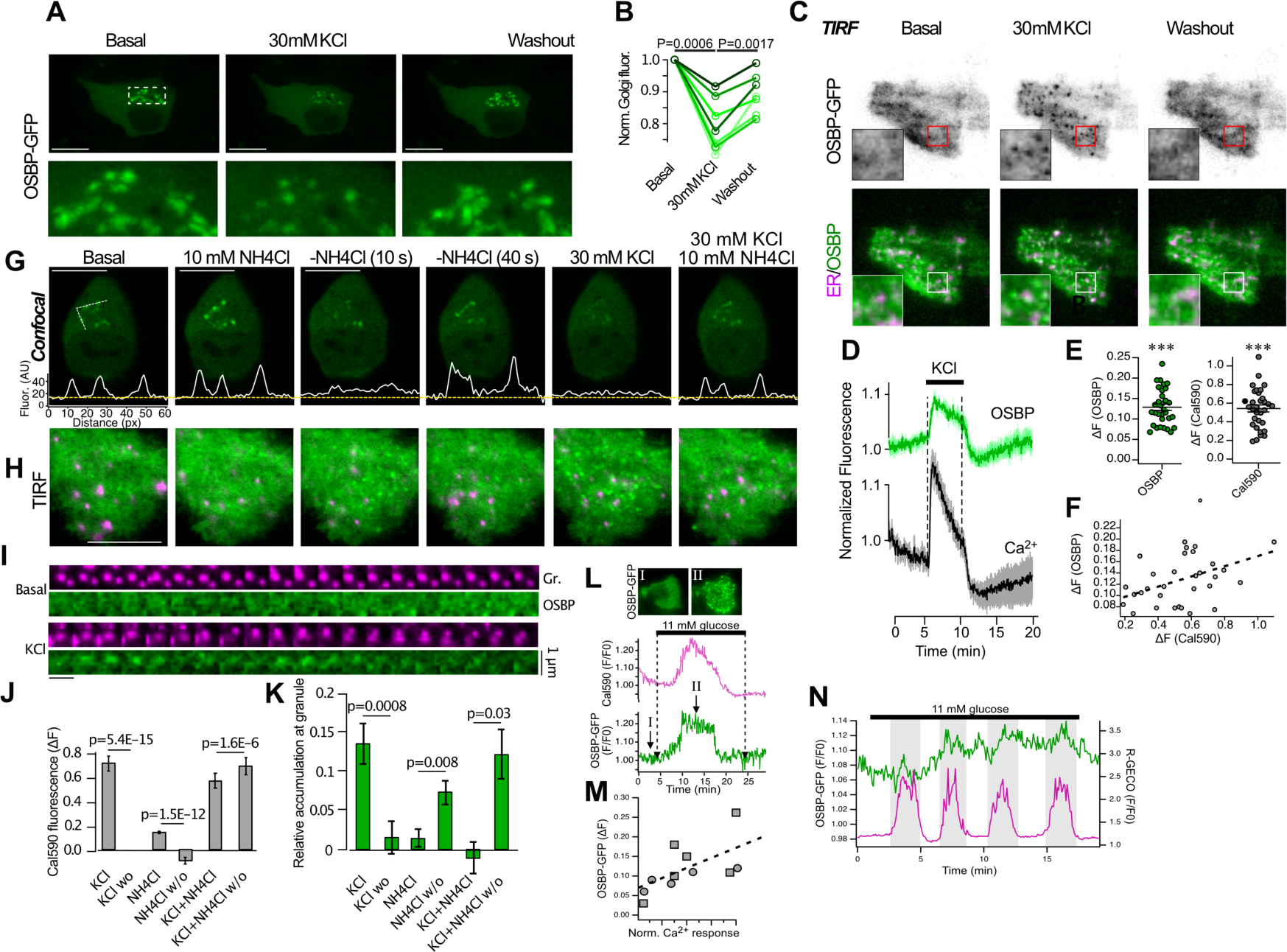
Spatial control of OSBP by Ca^2+^ and cytosolic pH. **A.** Confocal microscopy images of a MIN6 cell expressing GFP-OSBP exposed to 30 mM KCl. Images below are magnifications from the boxed area. **B.** Normalized Golgi OSBP-GFP fluorescence before, during and after stimulation with 30 mM KCl (n=7 cells; 2-tailed paired Student’s t-test). **C.** TIRF microscopy images of a MIN6 cell expressing GFP-OSBP (green) and ER-mRFP (magenta) before, during and after depolarization with 30 mM KCl. Notice the appearance of GFP-OSBP puncta that overlap with the ER-marker following depolarization. **D.** TIRF microscopy recordings from MIN6 cells expressing OSBP-GFP (green) and loaded with the Ca^2+^ indicator Cal590 (means±S.E.M. for 32 cells from 3 experiments). **E.** Quantification of the plasma membrane fluorescence change in response to depolarization in MIN6 cells expressing GFP-OSBP and loaded with Cal590 (*** P<0.001 for comparison to 0; n=32 cells from 3 experiments). **F.** Correlation between the depolarization-induced increase in Ca^2+^ and the appearance of GFP-OSBP at the plasma membrane. **G.** Confocal microscopy images of a MIN6 cell expressing GFP-OSBP and exposed to the indicated solutions. White line is an intensity profile plot across the Golgi. **H.** TIRF microscopy images of a MIN6 cells expressing GFP-OSBP (green) and NPY-mCherry (magenta) during stimulation with the indicated solutions. Notice the appearance of small, punctate structures following depolarization and acidification. **I.** Cut-outs showing the dynamic localization of GFP-OSBP to NPY-mCherry-positive granules (Gr.) following depolarization with 30 mM KCl. **J.** Quantification of Cal590 fluorescence change in response to the indicated stimulations. **K.** Quantifications of the relative accumulation of GFP-OSBP at NPY-mCherry positive granules in response to the indicated stimulations. **L.** TIRF microscopy recording of GFP-OSBP (green) and Cal590 (magenta) fluorescence during exposure to 11 mM glucose. Images from before and during glucose stimulation are shown above. **M.** Correlation between the glucose-induced increase in Ca^2+^ and the appearance of GFP-OSBP at the plasma membrane. **N.** TIRF microscopy recording of GFP-OSBP and R-GECO from a MIN6 cell during exposure to 11 mM glucose. Notice that Ca^2+^ oscillations are mirrored by increases in GFP-OSBP at the plasma membrane.

### Coordination of OSBP and Sac2 on the surface of SGs controls insulin secretion

Both PI(4)P and cholesterol are important components of insulin SGs that contribute to normal granule release. We and others recently showed that the PI(4)P phosphatase Sac2 localized to SGs (Cao et al., 2020; Nguyen et al., 2019), and that loss of Sac2 results in excess accumulation of SG PI(4)P. Insulin SGs are also a major storage site for cellular cholesterol (Galli et al., 2023), shown here by the strong colocalization between the fluorescent cholesterol marker filipin and the SG marker NPY-GFP (Fig. 6A). To explore the potential connection between SG cholesterol and PI(4)P, we performed siRNA-mediated knockdown of Sac2 in MIN6 cells and visualized cellular cholesterol using both fluorescently label cholesterol (Bodipy-cholesterol) and Filipin. We found that Sac2 knockdown resulted in strong accumulation of cholesterol on intracellular structures that colocalized with a marker of insulin SG (Fig. 6C-F), indicating that SG PI(4)P may participate in cholesterol counter-transport. To test whether this transport depends on OSBP, we reduced OSBP expression by siRNA knockdown. OSBP knockdown alone caused a slight reduction in intracellular cholesterol accumulation, but, importantly, simultaneous knockdown of OSBP and Sac2 prevented the excess SG cholesterol accumulation induced by Sac2 knockdown (Fig. 6G-I). These results indicate an interdependence of Sac2 and OSBP in controlling SG PI(4)P and cholesterol levels.

**Figure 6.**
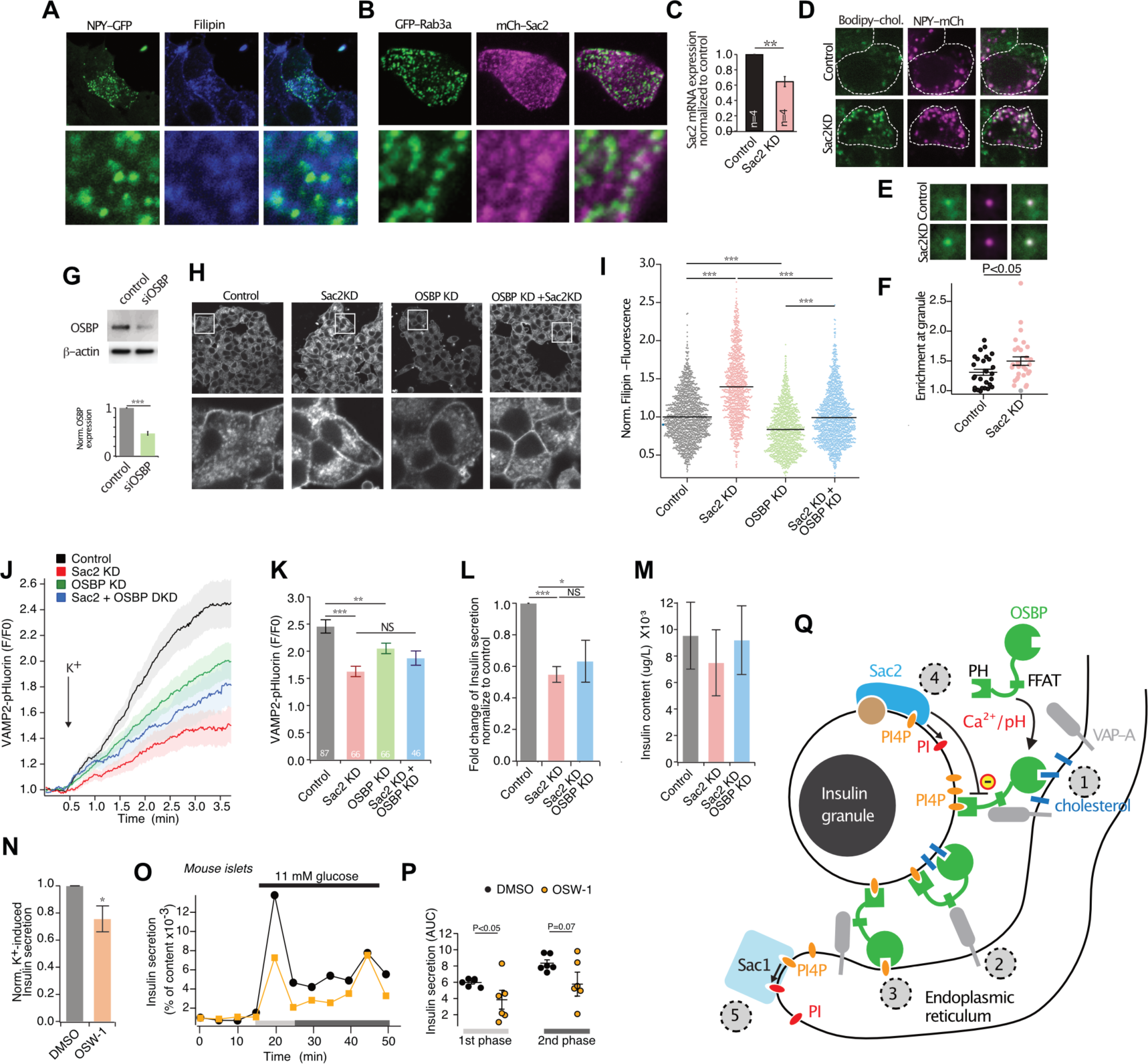
OSBP and Sac2 coordinately regulate insulin secretion. **A.** Immunofluorescence image of MIN6 cells expressing NPY-GFP (green) and stained with Filipin (blue) to visualize cholesterol. **B.** Confocal microscopy images of a MIN6 cells expressing the granule marker GFP-Rab3a (green) and mCherry-Sac2 (magenta). **C.** Quantitative RT-PCR determination of Sac2 mRNA levels in control and Sac2 knockdown cells expressed relative to control after normalization to GAPDH mRNA levels (n=4, ** P<0.01). **D.** Confocal microscopy images of NPY-mCherry-expressing MIN6 cells loaded with fluorescent cholesterol (Bodipy-cholesterol). Top panel shows cells transfected with control siRNA and bottom panel shows cells transfected with siRNA against Sac2. **E.** Confocal microscopy images showing the average accumulation of Bodipy-cholesterol fluorescence at NPY-mCherry positive structures in control cells or Sac2 knockdown cells. Images are averages of 270 and 290 structures from 27 and 29 cells. **F.** Scatter plot showing the enrichment of bodipy-cholesterol at NPY-mCherry positive structures (n=27 cells for control and 29 cells for Sac2 KD; 2-tailed unpaired Student’s t-test). **G.** Example immunoblot of lysates from MIN6 cells treated with control or OSBP siRNA probes with antibodies against OSBP and β-actin along with quantifications of OSBP protein levels in MIN6 cells treated with control or OSBP siRNA (expressed relative to β-actin; n=3; *** P<0.001, Mann-Whitney U-test). **H.** Confocal microscopy images of filipin-stained MIN6 cells treated with control siRNA, siRNA against Sac2, OSBP or Sac2+OSBP. **I.** Normalized cytoplasmic Filipin intensity in control MIN6 cells (n=1604) or MIN6 cells treated with siRNA against Sac2 (n=1176), OSBP (n=1266) or Sac2+OSBP (n=1492) (*** P<0.0001; ANOVA followed by Tukey’s post hoc test). **J.** TIRF microscopy recordings of VAMP2-pHluorin fluorescence change in response to 30 mM K^+^ in control cells and cells treated with siRNA against Sac2, OSBP or Sac2+OSBP. Data presented as means±S.E.M. for 47 (control), 31 (sac2 KD), 42 (OSBP KD) and 21 (Sac2+OSBP KD) cells from one experiment. **K.** Means±S.E.M. for the maximum VAMP2-pHluorin fluorescence increase in response to 30 mM K^+^ in the indicated cells. Data from 87 (control), 66 (Sac2 KD), 66 (OSBP KD) and 46 (Sac2+OSBP KD) cells from three experiments (** P<0.01; *** P<0.001, ANOVA followed by Kruskall-Wallis post hoc test). **L.** ELISA measurements of glucose-stimulated insulin secretion from MIN6 cells treated with control siRNA or siRNA against Sac2 or Sac2+OSBP. Data is expressed as fold change in insulin secretion when the glucose concentration was increased from 3 mM to 20 mM and is expressed relative to control cells (averages±S.E.M.; n=5; * P<0.05; *** P<0.001; Mann-Whitney U-test). **M.** Insulin content in control cells or cells treated with control siRNA or siRNA against Sac2 or Sac2+OSBP (averages±S.E.M.; n=5). N. ELISA measurement of K^+^-induced insulin secretion in DMSO-(control) and OSW-1-(20 nM, 60 mins) treated MIN6 cells (n=4; * P<0.05; Mann-Whitney U-test). **O.** Insulin secretion from mouse islets perfused with the indicated glucose buffers. The islets were pretreated for 30 mins with DMSO (0.1 vol%; black) or OSW-1 (20 nM, orange). **P.** Quantification of the first and second phase insulin secretion from the perfused mouse islets in K (n=5 for each condition; Mann-Whitney U-test). **Q.** Cartoon illustrating the cooperation between granule-localized Sac2 and OSBP in controlling turnover of granular PI(4)P and counter-transport of cholesterol. 1) OSBP binds to the ER via interactions between the FFAT-motif and VAP-A and to the insulin granule via interactions between the PH-domain and PI(4)P. OSBP extracts a cholesterol lipid from the ER membrane that is harbored within the ORD domain. 2) The cholesterol lipid is delivered to the insulin granule membrane. 3) A PI(4)P lipid is extracted from the granule membrane and delivered to the ER membrane. 4) The association of OSBP with the insulin granule is negatively regulated by the PI(4)P phosphatase Sac2. 5) PI(4)P delivered to the ER membrane is rapidly dephosphorylated by the PI(4)P phosphatase Sac1.

To test if the Sac2 and OSBP-dependent changes in intracellular cholesterol had an impact on insulin secretion, we determined the effect of Sac2 or OSBP knockdown on depolarization-induced insulin secretion using an optical assay based on VAMP2-pHluorin and TIRF microscopy. Consistent with previous studies, both Sac2 KD and OSBP KD resulted in reduced insulin secretion (34% and 16%, respectively; Fig 6J,K) (Nguyen et al., 2019). Simultaneous Sac2 and OSBP knockdown suppressed secretion to the same extent as OSBP knockdown alone (24%), and secretion was still significantly lower than under control conditions (Fig. 6J, K). Similar results were obtained when glucose-stimulated insulin secretion was assessed using ELISA. siRNA-mediated knockdown of Sac2 reduced the secretory response to 55% of control (P<0.001), and simultaneous knockdown of OSBP did not restore the response to the level of control cells (Fig. 6L). Insulin content was unaffected by both Sac2 and Sac2/OSBP knockdown, indicating that the reduction in insulin secretion is due to impaired release and not production (Fig. 6M). To more acutely test the involvement of OSBP in the regulation of insulin secretion, we incubated MIN6 cells with OSW-1 (20 nM) for 30 min and assessed the effect on depolarization-induced insulin secretion by ELISA. This timepoint was chosen because it is shorter than the time required for de novo synthesis of insulin SGs at the TGN (Davidson et al., 1988), ruling out potential effects of OSW-1 treatment on SG biogenesis. We found that OSBP inhibition suppressed insulin secretion by 26±9% (Fig. 6N) but was without effect on the voltage-dependent Ca^2+^ influx (Suppl. Fig. 6). OSW-1 also inhibited glucose-stimulated insulin secretion from isolated mouse pancreatic islets but was without effect on glucose-induced Ca^2+^ influx in the same islet preparation (Fig. 6O, P and Suppl. Fig. 6). These results show that lipid transport by OSBP is involved in the acute regulation of insulin secretion downstream of SG formation at the Golgi. The findings are consistent with a cooperative action between Sac2 and OSBP on the surface of SGs of importance for normal insulin secretion (Fig. 6Q).

## DISCUSSION

In this study we find that the increase in SG PI(4)P that occurs as a consequence of reduced Sac2 expression (Nguyen et al., 2019) is paralleled by the accumulation of cholesterol in the same compartment. This accumulation is caused by recruitment of the PI(4)P/cholesterol exchange protein OSBP to the surface of SGs, where it participates in the formation of ER-SG contact sites through interactions with ER-localized VAP proteins and PI(4)P on the granule surface. siRNA-mediated knockdown of Sac2 or OSBP, as well as pharmacological inhibition of OSBP, all resulted in impaired insulin secretion. Together, these results show for the first time the existence of physical contacts between the ER and SGs and demonstrate that these are sites of lipid transfer of importance for normal insulin secretion.

The cholesterol concentration in the ER is only around 5 mol% of membrane lipids (Van Meer et al., 2008), but it increases in the Golgi, at least in part through the action of OSBP. Mature insulin SGs are estimated to contain ∼35 mol% cholesterol (Galli et al., 2023; Tsuchiya et al., 2010), comparable to concentrations found in the plasma membrane (Van Meer et al., 2008). Given that a typical β-cell contains around 10,000 SGs with a total surface area 4.5 times larger than the plasma membrane (Sato Tamiko, 1981), the granule pool of cholesterol represents a major storage site of the lipid in β-cells. The importance of cholesterol for normal insulin secretion has been extensively studied. For example, cholesterol depletion alters glucose metabolism and interferes with the localization of ion channels and SNARE proteins, leading to aberrant insulin secretion regulation (Tsuchiya et al., 2010; Vikman et al., 2009), whereas cholesterol overloading leads to docking and exocytosis defects (Bogan et al., 2012; Hao et al., 2007). Cholesterol transport within β-cells is also important for normal insulin secretion. Both ABCG1 and ABCA1 cholesterol transporters are involved in SG biogenesis, and loss of these transporters leads to cholesterol accumulation and impaired insulin secretion, at least in part by destabilizing the SGs, resulting in their lysosomal degradation (Hussain et al., 2018; Kruit et al., 2010). The ABC cholesterol transporters do not localize to SGs and it is generally assumed that cholesterol loading occurs prior to granule departure from the trans-Golgi network (Harris et al., 2018). OSBP, another cholesterol transport protein found at membrane contact sites, has also been shown to positively regulate insulin secretion, but its cellular localization in β-cell is not known (Hussain et al., 2018). We now confirm that loss of OSBP activity results in reduced insulin secretion. We also find that OSBP associates with a fraction of SGs via an interaction between its PH-domain and PI(4)P on the granule surface (Levine and Munro, 2002). The phosphatase Sac2 negatively regulates PI(4)P levels on SGs (Nguyen et al., 2019), and knockdown of Sac2 resulted in accumulation of granule PI(4)P and in excess cholesterol loading that could be normalized by simultaneous OSBP knockdown. Acute inhibition of OSBP with OSW-1 (Burgett et al., 2011) resulted in redistribution of OSBP to SGs, which depended both on an intact C-terminal PH-domain and on ER-localized VAP-A/B. OSBP is not strongly attached to target membranes, and efficient lipid transport depends on dynamic equilibrium between membrane-bound and free molecules and arises, as least in part, as a consequence of OSBP extracting the same lipid (PI[4]P) that it uses for membrane tethering (Mesmin et al., 2013; Mesmin et al., 2017). When the exchange of lipids is prevented by OSW-1, this results in stabilization of OSBP at PI(4)P-rich membrane structures, in this case insulin SGs. Lowering of granular PI(4)P levels, either by recruitment of Sac2 or inhibition of PI4-kinase, therefore destabilizes the interaction between OSBP and the SG membrane. The negative regulation of OSBP localization by Sac2 may be a way to avoid cholesterol overloading of SGs. Given that both cholesterol excess and deficiency cause defects in insulin secretion (Bogan et al., 2012; Hao et al., 2007; Vikman et al., 2009; Xia et al., 2008), such negative feedback may be of importance to ensure that SGs accumulate appropriate amounts of cholesterol. However, Sac2 likely regulates insulin secretion also distal to cholesterol loading, since simultaneous Sac2 and OSBP knockdown normalized SG cholesterol content but not insulin secretion. It is possible that other lipid transport proteins operate at ER-SG contact sites in a PI(4)P dependent manner. In fact, most ORPs have PI(4)P as the preferred binding target (Nakatsu and Kawasaki, 2021). One candidate is ORP3, which similarly to OSBP is equipped with a PI(4)P-binding PH-domain and a VAP-A-binding FFAT-motif. This protein has been shown to counter-transport PI(4)P and phosphatidylcholine (PC) at ER-PM contact sites (D’souza et al., 2020). SGs contain PC (MacDonald et al., 2015), but it is not known how the levels of this lipid are regulated at this cellular location or what the downstream effectors are. An attractive candidate is PLD1, which catalyzes the formation of phosphatidic acid from PC, localizes to SGs and is required for insulin secretion (Hughes et al., 2004; Ma et al., 2010). It is also possible that the changes in ER-SG contacts brought about by PI(4)P accumulation result in aberrant recruitment of other proteins to these sites. It would for example be of relevance to determine if contacts between ER-localized STIM1 and granule-localized Orai1 can form in response to changes in ER Ca^2+^ and contribute to cellular Ca^2+^ homeostasis, similar to what has been shown in other secretory cells (Dickson et al., 2012). Insulin-containing SGs represent a significant Ca^2+^ store in β-cells that also contribute to insulin secretion, although through what mechanisms remains unknown (Idevall-Hagren and Tengholm, 2020; Mitchell et al., 2001).

OSBP is primarily engaged in lipid transport at the ER-Golgi interface under resting conditions, yet OSBP has been show to participate in similar reactions at other membrane contact sites under different physiological and pathophysiological conditions (Nakatsu and Kawasaki, 2021). Mechanisms that control OSBP redistribution and dynamics are still poorly understood, but it has been shown that Ca^2+^ can displace OSBP from the Golgi (Balla et al., 2008; Malek et al., 2021). We now show that elevation of the cytosolic Ca^2+^ concentration, together with the ensuing acidification, resulted in weak displacement of OSBP from the Golgi but also in enhanced association between OSBP and SGs, at least in the sub-plasma membrane space of the β-cell. This is reminiscent of OSBP inhibition with OSW-1, which stabilizes the interaction between OSBP and SGs. How Ca^2+^ stabilizes OSBP at SGs and whether it is coupled to displacement of OSBP from the Golgi is not clear. Displacement of OSBP from the Golgi also occurred following inhibition of PI4-kinases, but this did not result in accumulation of OSBP at SGs, indicating either that PI4-kinase are required to maintain granule PI(4)P levels or that Ca^2+^/pH promote the association trough an additional mechanism. Such mechanism may involve pH-mediated changes in PI(4)P protonation (Shin et al., 2020), direct effects of Ca^2+^ on OSBP (Malek et al., 2021) or PI(4)P (Kang et al., 2017) as well as pH or Ca^2+^ induced clustering of PI(4)P in the target membrane (Redfern and Gericke, 2004; Sarmento et al., 2014). Irrespective of mechanism, the spatial control of OSBP by Ca^2+^ couples lipid exchange at the SG surface to Ca^2+^-triggered exocytosis of SGs. Consistently, we find that acute inhibition of OSBP activity suppresses insulin secretion, indicating that OSBP acts immediately upstream of the exocytic process and distal to the formation of SGs at the Golgi. Further support for this notion comes from our observation that both ER-SG contacts and OSBP binding preferentially involves SGs located close to the plasma membrane. These physically docked granules are preferentially released in response to depolarization or during the first phase of glucose-stimulated insulin secretion, and both these mechanisms are also suppressed by OSBP inhibition. In addition to being physically docked at the plasma membrane, these SGs also represent a subpopulation of granules that has recently been formed at the trans-Golgi network (Ivanova et al., 2013). It is therefore tempting to speculate that cholesterol transport at ER-SG contacts represents a key step in the maturation of newly formed SGs. Consistent with this notion, it has been shown that newly formed post Golgi secretory vesicles are enriched for ergosterol and have a higher membrane order than the late Golgi membranes (Klemm et al., 2009).

To conclude, we propose a model where glucose-induced increases in Ca^2+^, together with the ensuing acidification, facilitate interactions between OSBP and SGs. OSBP then participates in the controlled loading of cholesterol to the SG membrane in cooperation with the granule-localized PI(4)P phosphatase Sac2 in a process that is required for normal insulin secretion. OSBP interacts with granules at ER-SG contact sites but does not seem to be involved in the formation of these contacts. Future research should aim at identifying protein complexes and biological processes that occur at these largely uncharacterized MCSs.

## ACKNOWLEDGEMENTS

We are grateful for the expert technical assistance provided by Dr. Beichen Xie, Ms. Cornelia Carleson and Mrs. Antje Thonig. OI was supported by grants from The Swedish Research Council (MH-2019-01456); The Novo-Nordisk Foundation (NNF19OC0055275), Åke Wibergs stiftelse (M18-0146), Diabetesfonden (DIA2018-332), Magnus Bergwalls stiftelse (2018-02752), Familjen Ernfors stiftelse, EFSD/Lilly and EXODIAB. MS was supported with funds from the German Center for Diabetes Research (DZD e.V.) by the German Ministry for Education and Research (BMBF) and from the Innovative Medicines Initiative 2 Joint Undertaking under grant agreement No 115881 (RHAPSODY) and the Swiss State Secretariat for Education’ Research and Innovation (SERI; 16.0097-2). AM was the recipient of a MeDDrive grant from the Carl Gustav Carus Faculty of Medicine at TU Dresden. LE received support from The Swedish Research Council (2019-01406). ATengholm received support from The Swedish Research Council (2021-02081), Swedish Diabetes Foundation, Diabetes Wellness Foundation, Family Ernfors Foundation, Nils Erik Holmsten’s Foundation, Novo-Nordisk Foundation and the Swedish national strategic grant initiative EXODIAB (Excellence of diabetes research in Sweden). ATo was supported by UKRI Medical Research Council grants MR/R010676/1 and MR/X021467/1 as well as by UKRI COVID-19 Grant Extension Allocation (coA), and further support from the EFSD, Diabetes UK, Eli Lilly, the Commonwealth, and the Integrated Biological Imaging Network (IBIN).

