## Supplementary figures for "OSBP-mediated PI(4)P-cholesterol exchange at endoplasmic reticulum-secretory granule contact sites controls insulin secretion"

***Supplemental Figure 1: Controls for proximity ligation assay.*** **A.** Confocal images of MIN6 cells immunostained against insulin (cyan), Sec61β (yellow), and Rab3 (red) with the same antibodies used for PLA in Figure 1. **B.** Confocal images of MIN6 cells immunostained against insulin (cyan), VAP-A yellow), and Rab3 (red) with the same antibodies used for PLA in Figure 1. **C.** Confocal images for the negative PLA control performed with Sec61β as the only primary, omitting anti-Rab3, post-immunostained against insulin (cyan). **D.** Confocal images for the negative PLA control (yellow; right) performed with VAP-A as the only primary, omitting anti-Rab3, post-immunostained against insulin (cyan; left).

***Supplemental Figure 2: Validation of contact site reporter and antibodies.*** **A.** Diagram of ER-plasma membrane contact site detection using the ddRFP reporter system. Confocal images from MIN6 cells expressing the ER marker ER-oxGFP, RA-Sec61 (ER), and GB-CaaX (plasma membrane). Images are taken from z-planes showing a cross-section of the cell and a section at the bottom near the coverslip. **B.** Diagram and confocal image of a MIN6 cell expressing RA-Sec61 and GB-Sec61 (positive control). **C.** Diagram and confocal image of a MIN6 cell expressing only RA-Sec61 (negative control). The right image is contrast enhanced to show the very weak intrinsic RA fluorescence. **D.** Confocal image of a MIN6 cell expressing GFP-Rab3 as a granule marker and RA-Sec61. **E.** Diagram showing principle for testing ddRFP detection distance. RA-CaaX is targeted to the plasma membrane, while GB is fused either to the amino or carboxy terminus of E-Syt3 to control its distance from the PM. Below are TIRF microscopy images of MIN6 cells expressing ER-oxGFP, RA-CaaX, and either version of GB. Note that PM-targeted RA is not boosted by GB at ER-PM contact sites when a 20 nm separation is enforced (top pair). Cell outlined in blue is not expressing RA-CaaX **F.** Confocal images and magnifications of MIN6 cell expressing NPY-mNG, fixed and immunostained against chromogranin A (ChrgA) and Rab3a.

***Supplemental Figure 3: Effect of OSW-1 on ER and Golgi markers.*** **A.** Confocal images of MIN6 cells expressing NPY-mNG as a granule marker and VAP-A-Halo, treated with 20 nM OSW-1 for 20 mins. The top row shows the cells before treatment, and the bottom row is the same cells after treatment. Scale bar: 10 µm. **B.** Quantification of VAP-A positive insulin granules before and after treatment with OSW-1 as in A. **C.** Relative enrichment of VAP-A on insulin granules before and after treatment with OSW-1 analyzed with the same dataset as B. **D.** Confocal images of MIN6 cells treated with 20 nM OSW-1 for 20 mins, fixed and immunostained against VAP-A or VAP-B (magenta) and insulin (cyan). Scale bars: 10 nm. **E.** Confocal microscopy images of mouse islets treated with 20 nM OSW-1 for 20 mins, fixed and immunostained against GolgB1 (magenta) and insulin (green). **F.** Confocal microscopy images of mouse islets treated with 20 nM OSW-1 for 20 mins, fixed and immunostained against calnexin (magenta) and insulin (green). **G.** Quantifications of Golgi size and fluorescence upon treatment with either 20 nM OSW-1 or vehicle (DMSO), obtained from the GolgB1 immunostaining in E.

***Supplemental Figure 4: Effect of PI4-kinase inhibitors on OSW-1 induced OSBP redistribution.*** **A.** Confocal images of MIN6 cells expressing NPY-mNG (green) and Halo-OSBP (magenta), imaged live in basal imaging buffer, after 20 mins with the indicated PI4K inhibitor (or vehicle), and after 20 mins with 20 nM OSW-1. Scale bars:10 nm.

***Supplemental Figure 5: Response of OSBP to cellular perturbations.*** **A.** TIRF microscopy images of a MIN6 cell expressing GFP-OSBP following the indicated stimulations. Images have been contrast enhanced to emphasize the granular OSBP structures. **B.** TIRF microscopy images of a small region of a MIN6 cell expressing GFP-OSBP (green) and NPY-mCherry (magenta) before addition and after removal of 20 mM $NH_4Cl$. Line profiles to the right indicate fluorescence intensities across the images. Notice the enrichment of OSBP on NPY-positive structures following acidification. **C.** TIRF microscopy recording of BCECF fluorescence from a MIN6 cell stimulated as indicated. **D.** TIRF microscopy recording of Cal590 fluorescence from a MIN6 cell stimulated as indicated. **E-G.** Confocal microscopy images of NPY-mNG (granules; green) and HaloJF646-OSBP (magenta) before and after addition of 100 µM carbachol (E), 30 mM KCl (F) or washout of 20 mM $NH_4Cl$ (G). Quantified below is the change in OSBP Golgi/cytosolic ratio in response to the different treatments (n=5-8 experiments; 2-tailed paired Student's t-test). **H.** TIRF microscopy recordings from MIN6 cells expressing cytosolic pHluorin and exposed to DMSO (0.1%) or 20 nM OSW-1 (addition indicated by arrow), followed after 20 min by exposure to 30 mM KCl. Notice that OSW-1 causes acidification of the cytosol but does not prevent the subsequent KCl-induced acidification. Data presented as means±S.E.M. from 89 (DMSO) and 102 (OSW-1) cells.

***Supplemental Figure 6: Effect of OSW-1 on intracellular $Ca^{2+}$ dynamics.*** **A.** Cal520 fluorescence change in response to 30 mM $K^+$ in DMSO- or OSW-1- (20 nM, 60 mins) treated cells (means±S.E.M.; n=100 cells for both conditions). **B.** Cal520 fluorescence change in response to 11 mM glucose in DMSO- or OSW-1- (20 nM, 30 mins) treated mouse islet β-cells (n=5 islets per condition). **C.** Example recordings of cytosolic $Ca^{2+}$ concentration changes in mouse islet cells in response to 20 mM glucose. The islets were preincubated in DMSO (black) or 20 nM OSW-1 (orange) for 30 mins.

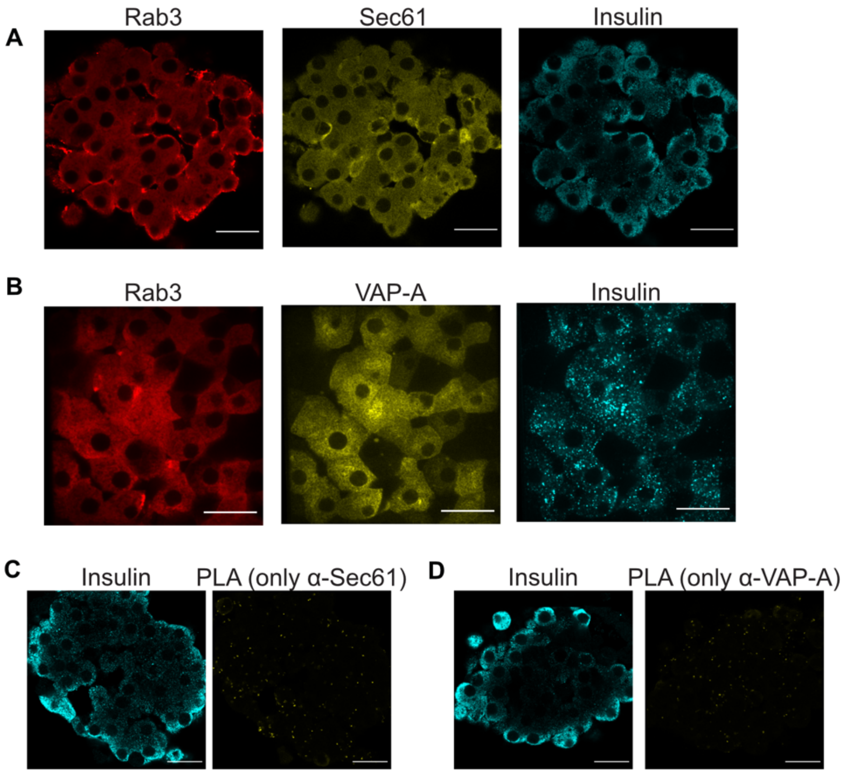


**Suppl. figure 1**

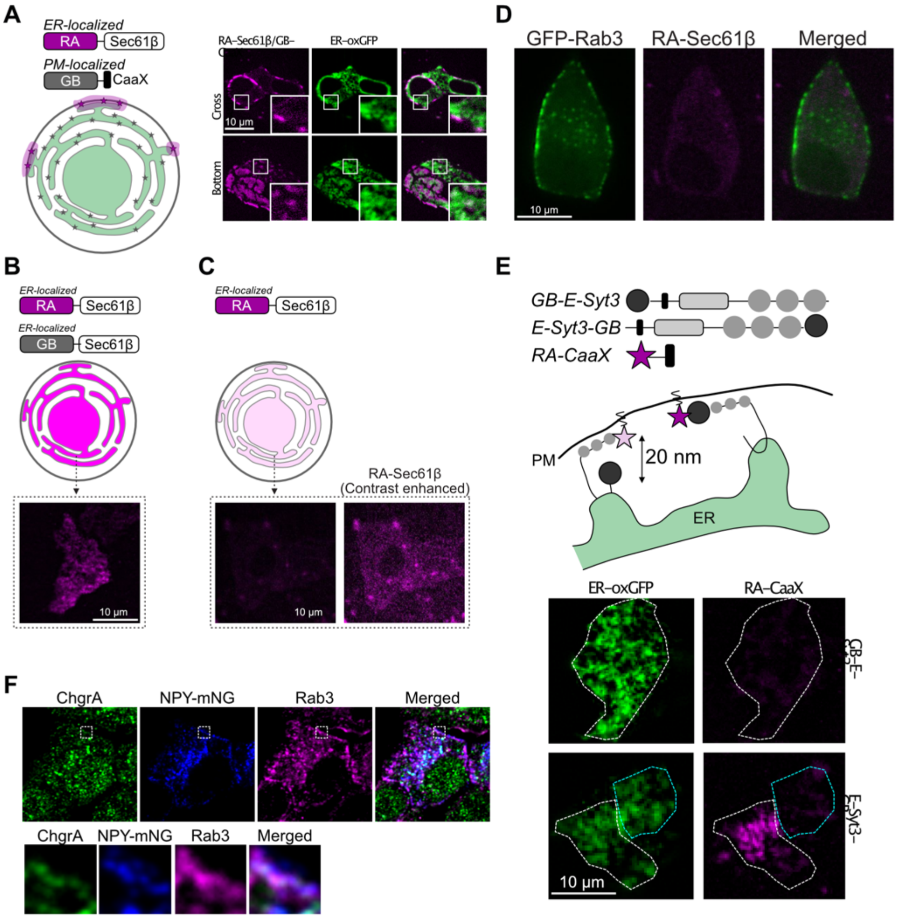


**Suppl. figure 2**
**(Related to Fig. 2)**

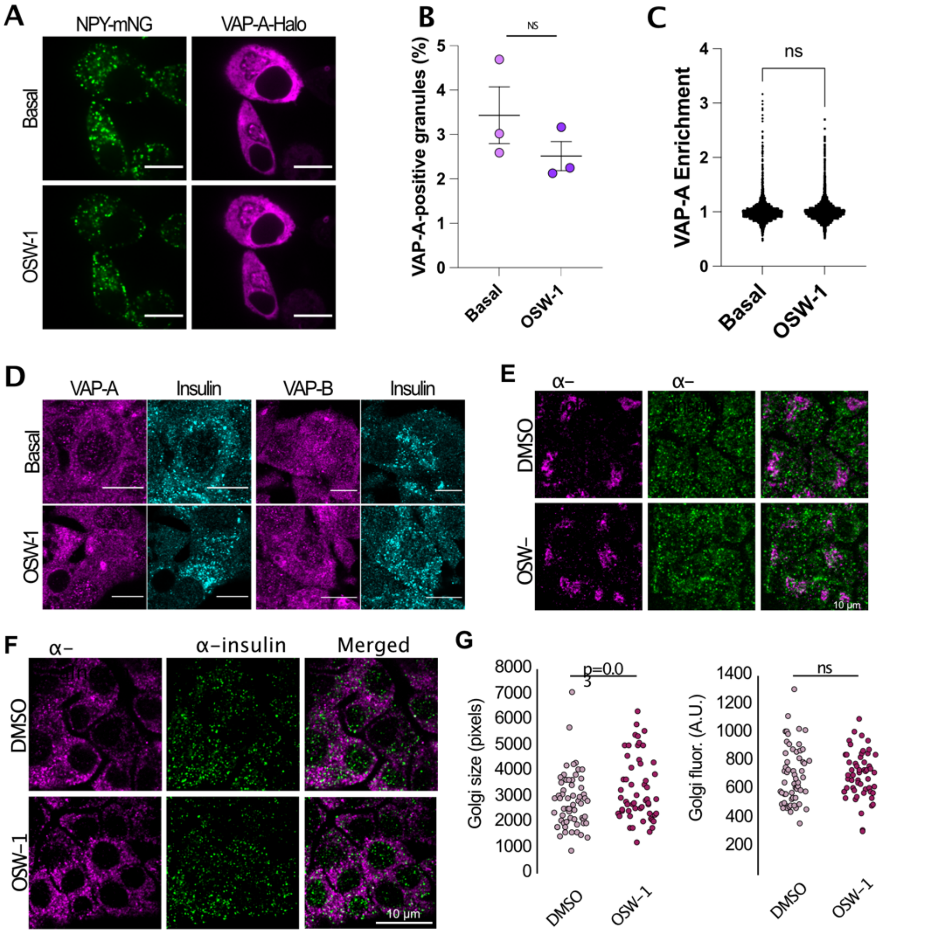


**Suppl. figure 3**

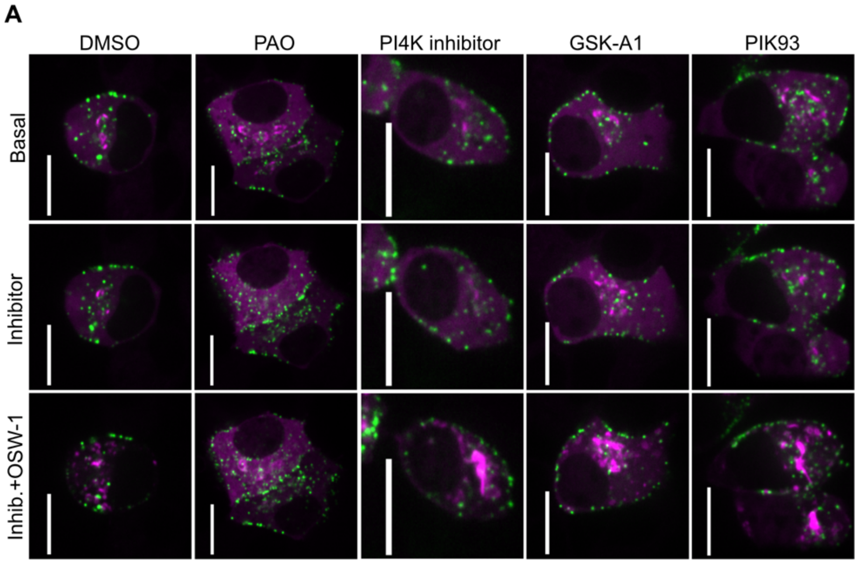


**Suppl. figure 4**
**(Related to Fig. 3)**

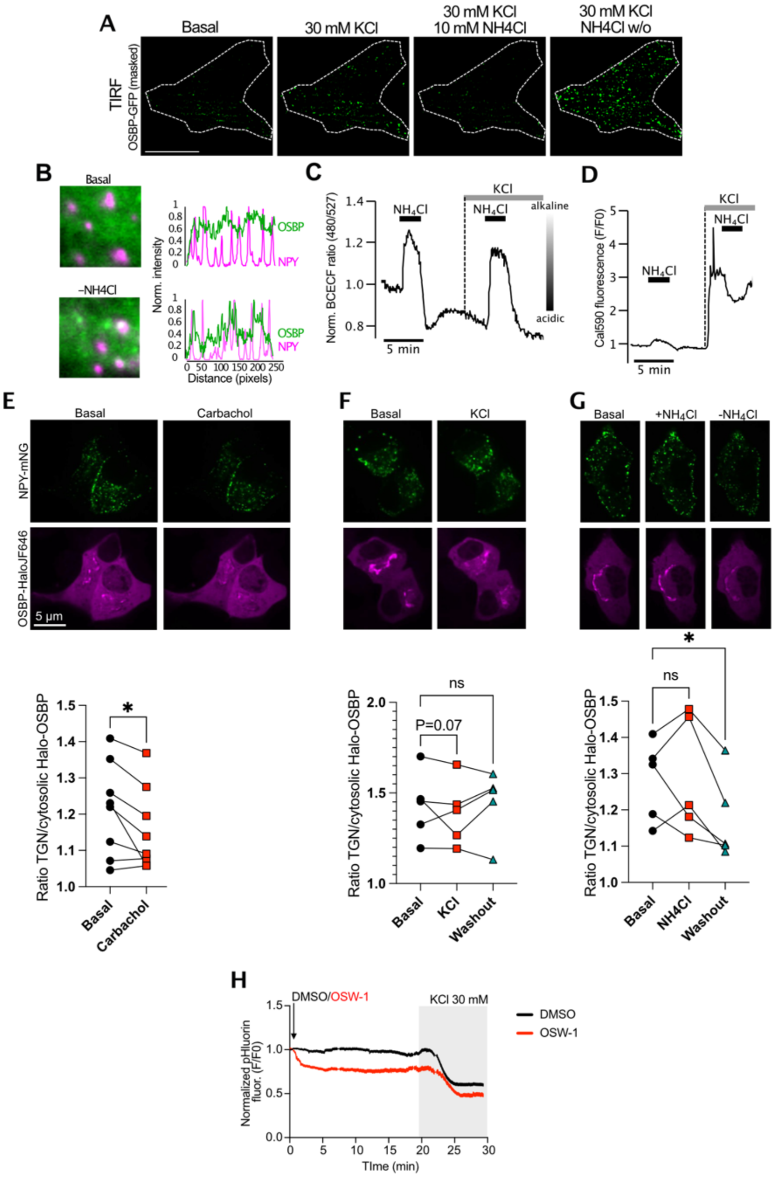


### Suppl. figure 5

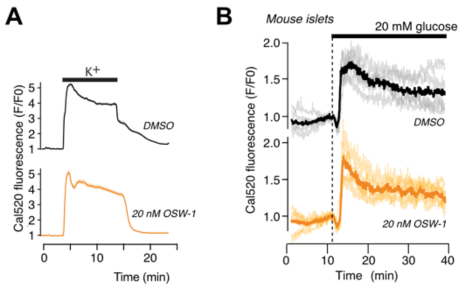


### Suppl. figure 6
